# Copper dependent ERK1/2 phosphorylation is essential for the viability of neurons and not glia

**DOI:** 10.1101/2021.03.18.435926

**Authors:** Kaustav Chakraborty, Sumanta Kar, Bhawana Rai, Reshma Bhagat, Nabanita Naskar, Pankaj Seth, Arnab Gupta, Ashima Bhattacharjee

**Author notes:** Correspondence: Ashima Bhattacharjee, Address: Amity University Kolkata, Plot no 36,37 &38, Major Arterial Road, Action area II, Kadampukur Village, Rajarhat, Newtown, West Bengal – 700135, India. Ph no: 7044095662. Department of Genetics, Washington University in Saint Louis, St Louis, MO, USA.

## Abstract

Intracellular Copper [Cu(I)] has been hypothesized to play role in the differentiation of the neurons. This necessitates understanding the role of Cu(I) not only in the neurons but also in the glia considering their anatomical proximity, contribution towards ion homeostasis, neuronal physiology, and neurodegeneration. In this study we did a systematic investigation of the changes in the cellular copper homeostasis during neuronal and glial differentiation and the pathways triggered by them. Our study demonstrates increased mRNA for the plasma membrane copper transporter CTR1 leading to an increased Cu(I) during neuronal (PC-12) differentiation. ATP7A is retained in the Trans Golgi Network (TGN) despite high Cu(I) demonstrating its utilization in triggering the pathways towards the neuronal differentiation. One of these pathways is ERK1/2 phosphorylation accompanying the differentiation of both PC-12 and human fetal brain derived neuronal progenitor cells. The study demonstrates that the ERK1/2 phosphorylation is essential for the viability of the neurons. In contrast, differentiated C-6 (glia) cells contain low intracellular copper and significant downregulation of the ERK1/2 phosphorylation. Interestingly ATP7A shows vesicular localization despite the low copper in the glia. In addition to the TGN in the perinuclear region, ATP7A localizes into RAB11 positive recycling endosomes in the glial neurites, not observed in the neurons. Our study demonstrates role of the copper dependent ERK1/2 phosphorylation in the neuronal differentiation. Whereas glial differentiation largely involves sequestration of Cu(I) into the endosomes potentially (i) for ready release to the neurons (ii) rendering cytosolic copper unavailable for pathways like the ERK1/2 activation.

## Introduction

Brain is the largest storehouse of copper, second to the liver. Copper level varies in different regions of the brain as well as at different stages of the development. Copper level in the subventricular zone of rat brain increases in an age dependent manner (1). However, copper level decrease with age in the human brain (2). Variation in the level, at the developmental stages, reflects changes in its utilization and its importance. Besides copper distribution also varies in the different regions of the brain (3) thereby reflecting the heterogeneity in its utilization. However, the exact nature of utilization of the Cu(I) towards triggering pathways during the neuronal and glial differentiation is not well understood. The critical role of the maintenance of the copper level in the CNS is emphasized by the occurrence of CNS dysfunctions associated with the copper homeostasis disorders. Mutation in the copper transporting ATPase ATP7B leads to accumulation of the excess intracellular copper in the liver, kidney and brain known as Wilson Disease (WD) (4). Apart from the hepatic symptoms, WD involves neuropsychiatric symptoms (5). Defects in ATP7A causes three X-linked recessive disorders – Menkes Disease, Occipital Horn Syndrome and Spinal Muscular Atrophy. Menkes Disease, caused by systemic copper deficiency, involves atrophy of gray and white matter involving focal degeneration of all layers of cerebral cortex extending to other regions of the CNS. The degeneration involves both the neurons and the astrocytes (6). The mo-br mice exhibit distinct neuronal and glial abnormalities suggesting role of the intracellular copper in both the cell types. The abnormal axonal extension and synaptogenesis in these mice demonstrates role of copper towards the neurite extension (7). Moreover, ATP7A expression has also been detected in the developing neurites during synaptogenesis suggesting role of copper mediated pathways in the neurodevelopment related to learning and memory (8). This emphasizes the necessity to investigate the role of the intracellular copper homeostasis in the neuronal and glial differentiation.

In this study, we have investigated the changes in the cellular copper homeostasis during the neuronal and glial differentiation using two cell-based model system. Neuronal differentiation accompanies increased mRNA of the plasma membrane copper importer CTR1 and increased intracellular copper. Utilization of this excess copper is evident from the retention of the primary copper ATPase ATP7A in the Trans Golgi Network (TGN). One of the specific pathways triggered by the intracellular copper is phosphorylation of ERK1/2 during the neuronal differentiation. On other hand, the differentiated glia has low intracellular copper and subsequent downregulation of the ERK1/2 phosphorylation. Moreover, contrast to the neuronal cells, ATP7A localizes into the RAB 11 positive vesicles along the neurite processes in the glia despite low intracellular copper.

## Results

### (i) Differentiation of the PC-12 cells (neuron) involves increased intracellular copper in contrast to the differentiation of the C-6 (glia) cells

PC-12 cells are plated on collagen. To induce differentiation, cells are treated with Nerve Growth factor (NGF) as described in the methods. The cells showed increase in the neurite length (Fig 1A) and increased expression of B-III Tubulin (Fig 2A). However, unchanged expression of the Glial Fibrillary Acidic Protein (GFAP) demonstrates the purity of the neuronal lineage (Fig 2A). C-6 cells were plated and treated with dbCAMP and Theophyllin for 48 hours, whereby they demonstrated progressive neurite growth and branching of the neurites (Fig 1A,B). Increased mRNA for GFAP and unchanged mRNA level for B-III tubulin confirms the glial differentiation (Fig 1B and 2B). The PC-12 derived neurons have increased intracellular copper compared to the undifferentiated ones (Fig 2C). Increased intracellular copper accompanying the neuronal differentiation suggests its importance towards the process. However, the glia, differentiated in serum deprived condition contain low intracellular copper (Fig 2D).

**Figure 1:**
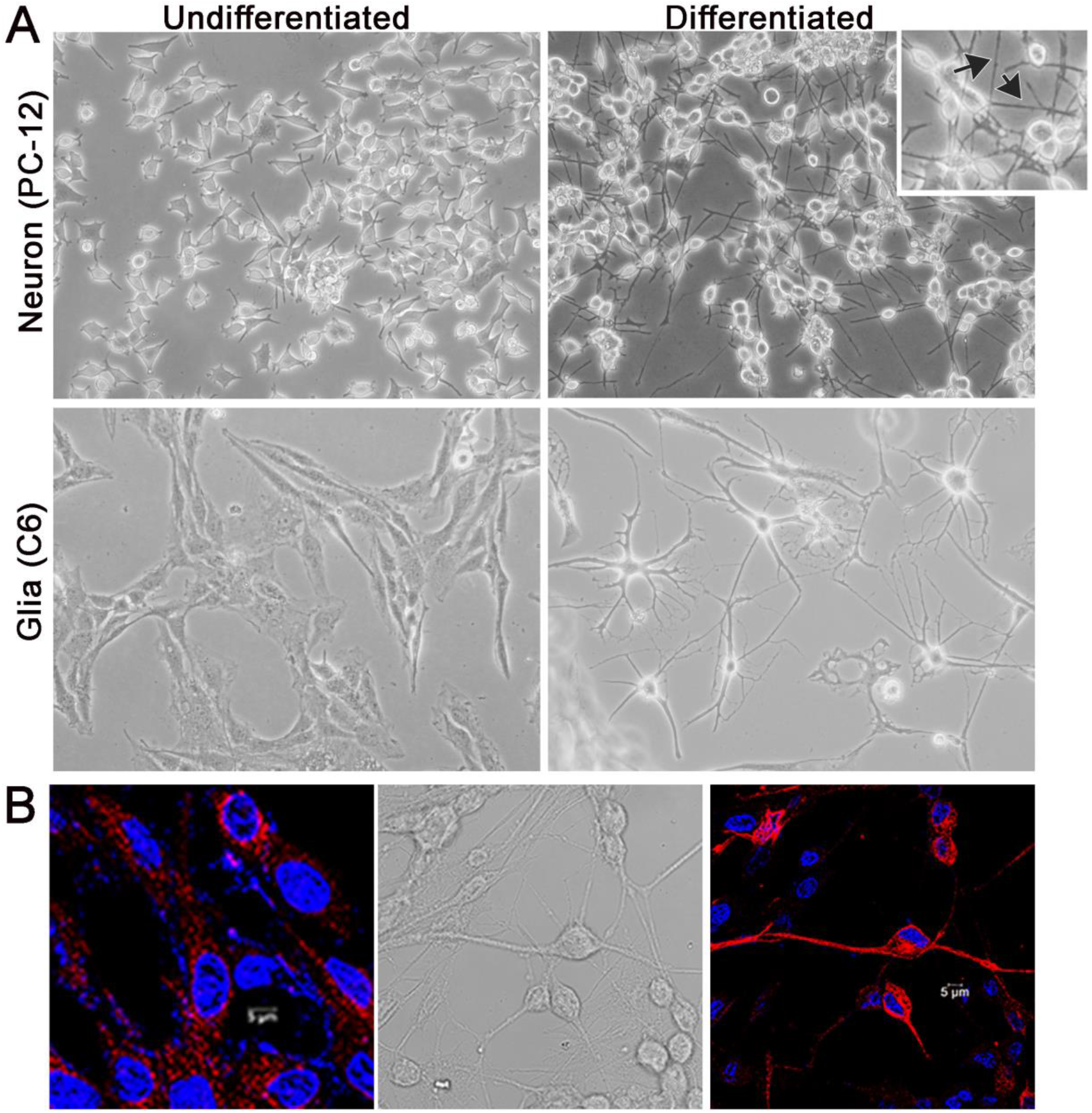
Differentiation of the PC12 and C6 cells into neuron& glia respectively. A (upper panel) - Bright field images (at 20X magnification) of undifferentiated and differentiated PC12cells. Increased neurite length is observed following treatment with Nerve Growth Factor (NGF). A (Lower Panel) - Bright field images (at 40X magnification) of undifferentiated and differentiated C6 cells. Increased length and branching of neurites are noticed with the treatment of dBcAMP and Theophylline. B– undifferentiated and differentiated C6 cells immunostained for GFAP. Increased expression of GFAP along the neurites confirms glial differentiation. Image acquired at 63X optical zoom, Scale bar, 5 μm.

**Figure 2:**
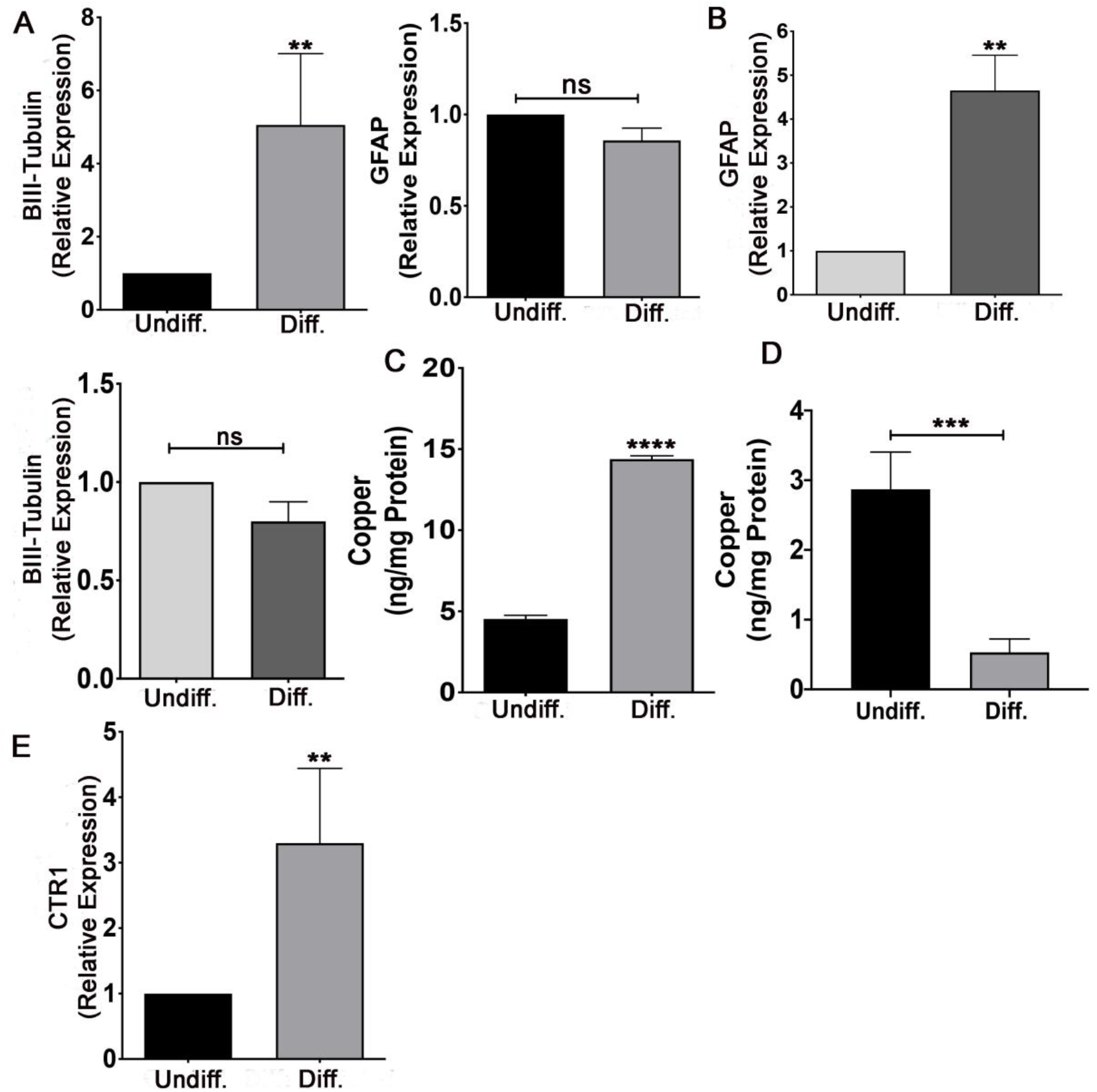
Changes in the cellular copper homeostasis accompanying the neuronal and the glial differentiation. A - Increased mRNA for β-III tubulin (left) and unchanged mRNA for GFAP (right) confirms differentiation of the PC-12 cells into the neurons (number of replicates - 4). B – Increased mRNA for GFAP and unchanged mRNA for B-III tubulin confirms differentiation of the C6 cells into the glia (number of replicates - 4). C - Level of the intracellular copper in the undifferentiated and the differentiated PC-12 cells as determined by ICP-MS. D – Level of intracellular copper in the undifferentiated and differentiated C-6 cells as determined by ICP-OES. E – Relative expression of the CTR1 mRNA in the undifferentiated and the differentiated PC-12 cells (number of replicates 3).

### (ii) Increased intracellular copper parallels increased expression of CTR1 mRNA in the differentiated PC-12 (neuron) cells but not in differentiated C-6 (glia) cells

To correlate how the intracellular copper, in the differentiated neurons, is distributed to the different intracellular locations, we investigated the mRNA expression of different chaperones and transporters designated to deliver copper to the different intracellular locations. Significant increase in the mRNA for CTR1 was observed (Fig 2E) in the differentiated PC-12 cells. This suggests that differentiation of PC-12 cells into the neurons involves an increased expression of the plasma membrane copper transporter CTR1 leading to the entry of copper into the cells. Moderate increase in CCS (Copper Chaperone for SOD1) and mitochondrial chaperone COX17 was observed (supplementary Fig1A) in the PC-12 derived neurons. This might be related to the increased mitochondrial copper demand accompanying the differentiation. Although intracellular copper is high, expression of mRNA for the primary copper ATPase-ATP7A and its delivery chaperone ATOX1 did not change (supplementary Fig 1A). Neither undifferentiated nor differentiated (PC-12) neurons and (C-6) glia expressed ATP7B (data not shown).

In contrast, Glial differentiation, does not show change in the mRNA expression for any chaperones and transporters including CTR1 (supplementary Fig 1B). Differentiation of C-6 cells into the glia is performed in copper deprived media (without serum) and the differentiated glia contain less intracellular copper (Fig 2D). Unchanged level of the CTR1 mRNA, under low intracellular copper, is intriguing. It suggests that maintenance of the low intracellular copper is crucial for the glial differentiation.

### (iii) ATP7A shows differential localization in the differentiated PC-12 (neuron) and C-6 (glia) cells

ATP7A traffics to the vesicles under high intracellular copper. Since the PC-12 derived neurons have high intracellular copper, we wanted to investigate if ATP7A shows vesicular localization. Interestingly, ATP7A showed localization exclusively on the TGN (Fig 3A) despite high intracellular copper. This suggests that the high intracellular copper in these neurons, is not excess and there might be utilization of the copper by multiple physiological pathways. Firstly, the neuronal copper, can be utilized towards the differentiation thereby retaining ATP7A in the TGN instead of the vesicles. Secondly, a bulk of the cytosolic copper can be transported into the Golgi based secretory pathway as already demonstrated (9) during the neuronal differentiation.

**Figure 3:**
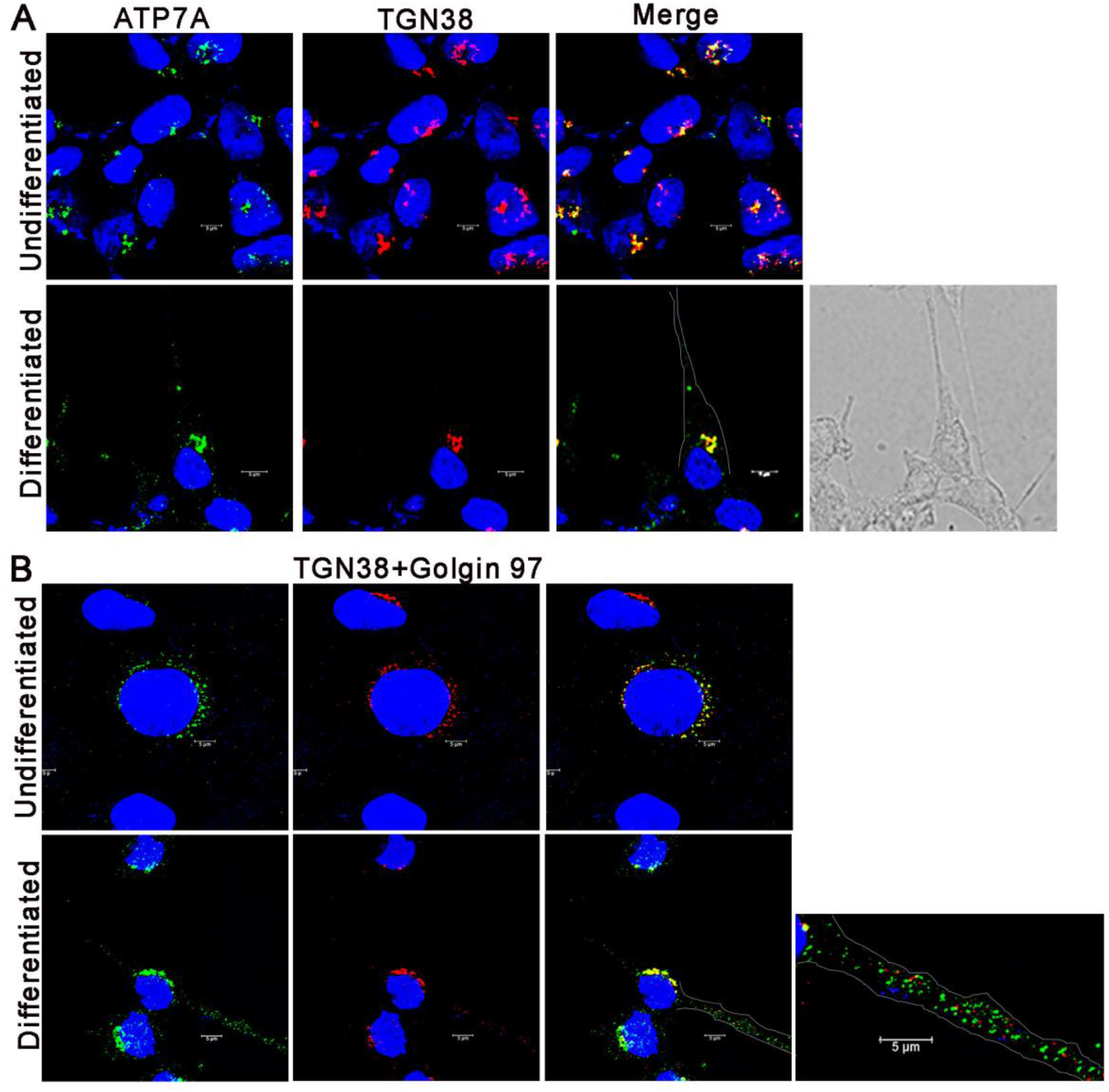
Localization of ATP7A in the undifferentiated and the differentiated PC-12 (neuron) and C-6 (glia) cells. A – Colocalization of ATP7A with TGN38 in the PC-12 cells. ATP7A colocalizes primarily with TGN38 in the perinuclear (soma) area. B – Colocalization of ATP7A with TGN38 and Golgin 97 in the C-6 cells. ATP7A localizes into the vesicular structures in the neurites in addition to the perinuclear localization. ATP7A colocalizes with TGN38 and Golgin97 in the perinuclear area. However, there is no colocalization of ATP7A containing vesicles in the neurites with the TGN markers.

In the differentiated C-6 glia, ATP7A shows colocalization with TGN38 and Golgin 97 in the perinuclear location. In addition, ATP7A also localizes into non-Golgi vesicular structures (Fig 3B, inset) along the neurites. Such vesicular localization of ATP7A is intriguing as the differentiated glia have low intracellular copper. TGN has been demonstrated to interact with the recycling endosomal system in the neurons playing important role in the trafficking of the synapse related proteins (9,10). Since role of copper has been implicated in the maturation of the synaptic vesicles and their release (11,12), we wanted to investigate if these ATP7A vesicles, in the glial neurites, could be TGN compartments.

To confirm this, we performed <u>S</u>timulated <u>E</u>mission <u>D</u>epletion <u>M</u>icroscopy (STED) to visualize these structures. At such resolution, ATP7A, in the vesicles is not seen to colocalize with TGN38 and Golgin 97 suggesting that these are not extensions of the Trans Golgi Network (TGN) (Supplementary Fig 2A). This suggests that ATP7A exits the TGN and localizes into the vesicles in the glia, despite low intracellular copper. Inorder to study the identity of these vesicles, we investigated if these vesicles are part of the canonical TGN exit pathway reported for ATP7B.

Lysosomal exocytosis has been demonstrated to be the mechanism of ATP7B mediated copper export (13). Although not demonstrated for ATP7A, we wanted to investigate if these vesicular structures play role in glial copper export involving the lysosomal route. We observed that the vesicular ATP7A, in the neurites do not colocalize with LAMP1 suggesting that these are not lysosomes (Supplementary Fig 3).

ATP7B has been reported to sort from the TGN to the basolateral endosomes prior to trafficking to the apical domain of the polarized WIF-B cells (14). It has been further demonstrated that MyosinVb localizes ATP7B to the gamma tubulin rich domains prior to apical sorting at high copper (15). Therefore, it is probable that post–TGN-exit both ATP7A and ATP7B follow a partially overlapping basolateral trafficking itinerary. The trafficking itinerary of ATP7A has not been studied in the polarized system. Since glia are unique polarized cells, we wanted to investigate if ATP7A vesicles are in the gamma tubulin rich domains. But vesicular ATP7A, in the glia, does not colocalize with gamma tubulin (Supplementary Fig 4) suggesting that the vesicles are not part of the Microtubule Organizing Centre (MTOC).

We further investigated if ATP7A, in these vesicles, localizes to the late endocytic pathway (16) for ready release of copper. STED microscopy showed no colocalization of ATP7A with Rab7 (Supplementary Fig 2B). Further we investigated if the ATP7A containing vesicular structures in the glial processes represent RAB11 (17) positive recycling endosomes. We observed colocalization of ATP7A and RAB11 in the vesicles along the glial (neurite) processes (Fig 4, Supplementary Fig 5). Additionally, we also observed considerable colocalization of ATP7A with RAB11 in the perinuclear region (Fig 4, Supplementary Fig 5) of both the differentiated and undifferentiated glial cells. This suggests that, under basal condition, ATP7A localizes to the recycling endosomes in addition to the TGN, in these cells. These RAB11 positive compartments extend to the neurites with differentiation and ATP7A resides into some of those compartments in the glia (Fig 4). Super-resolution microscopy (STED) shows ATP7A either in colocalization or in juxtaposition with RAB11 both in the perinuclear (Fig 5A) and along the neurites in the glia (Fig 5B).

**Figure 4:**
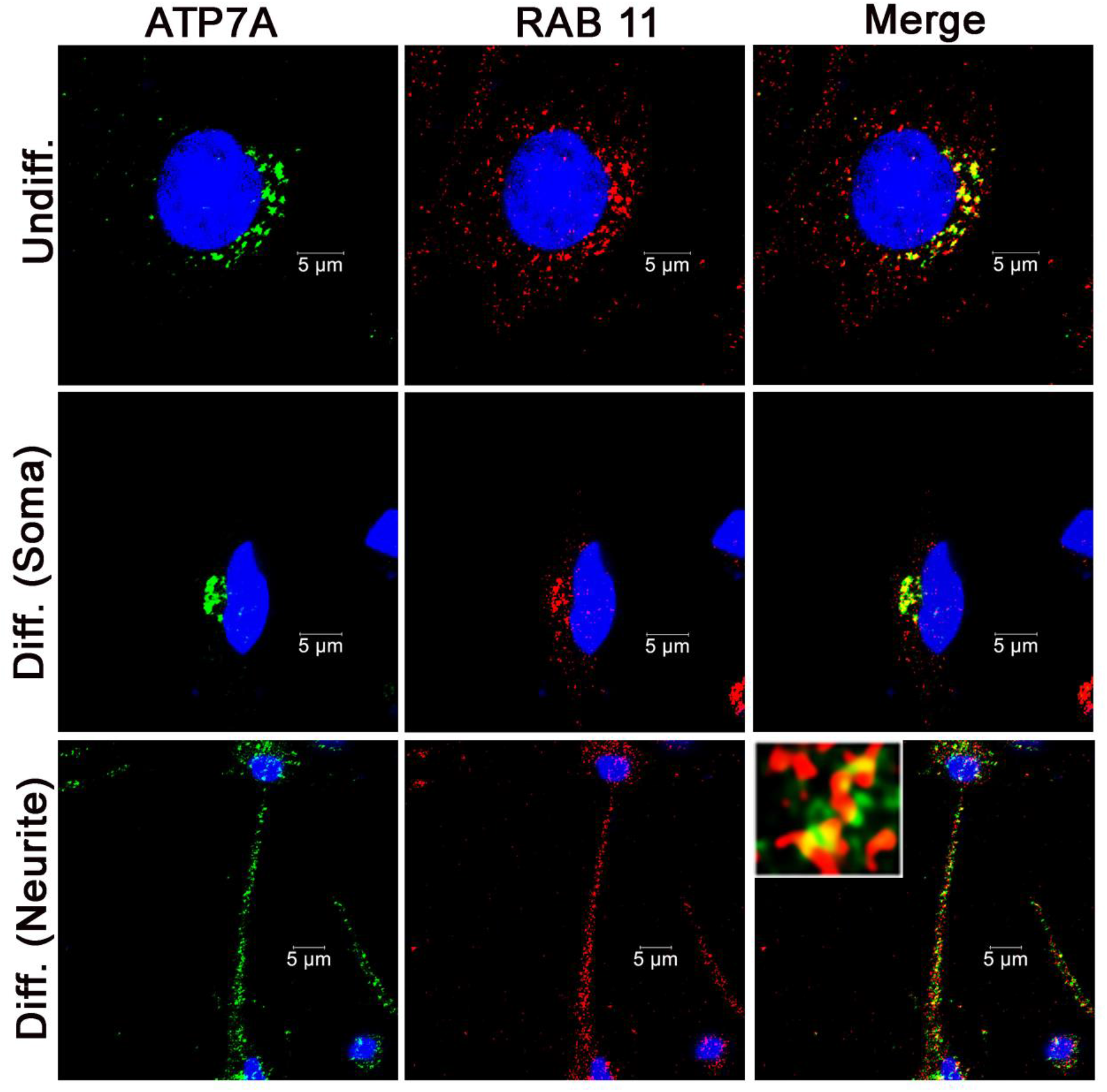
Localization of ATP7A in the RAB11 positive compartment in the perinuclear region (soma) and along the neurites in the differentiated C-6 (glia) cells.

**Figure 5:**
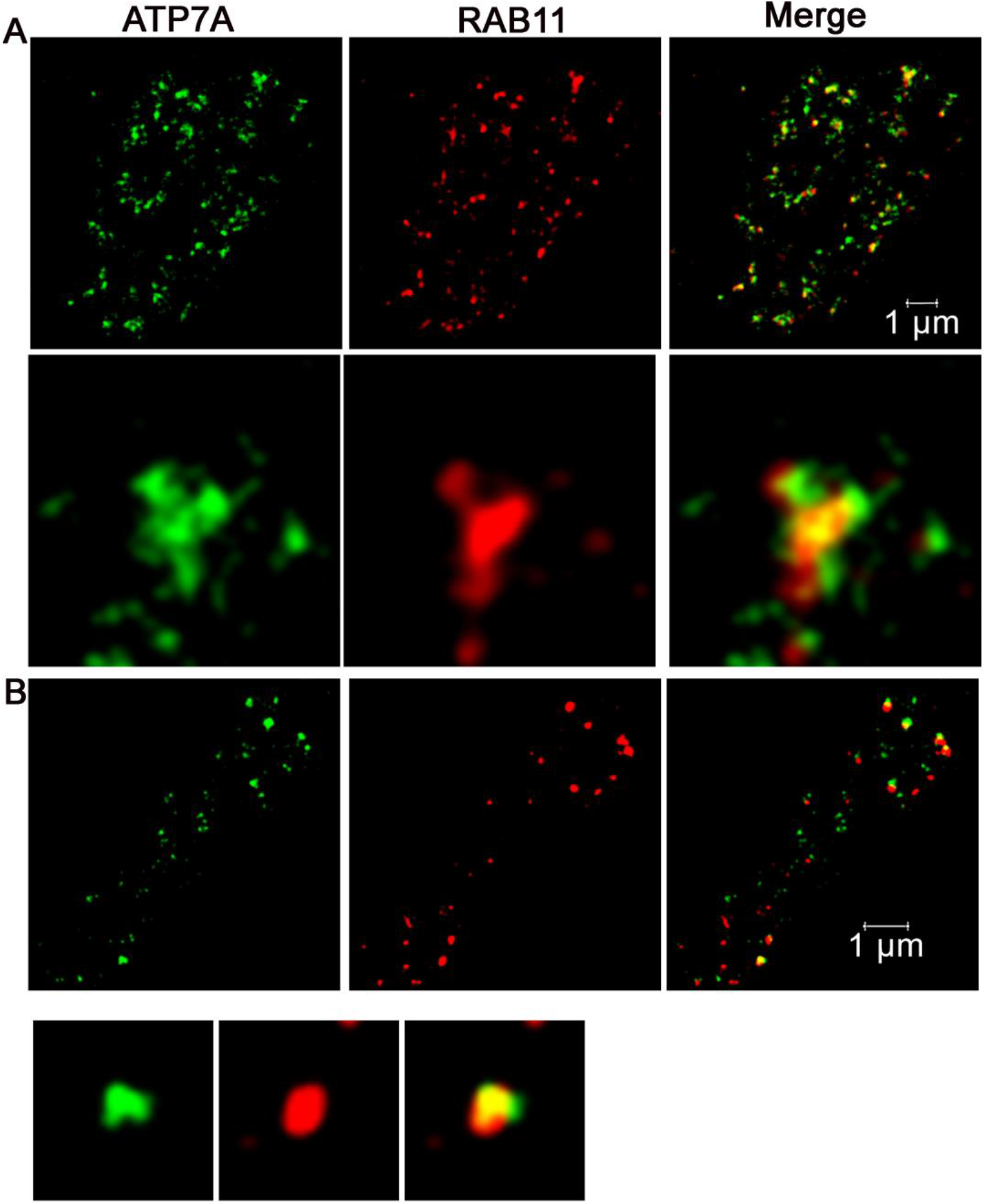
Colocalization of the ATP7A with RAB11 in the perinuclear (soma) and neurite in differentiated (C-6) glia as seen in STED microscopy. A (upper): ATP7A colocalizes with RAB11 in the perinuclear soma. A (lower): A Magnified area from the upper panel shows both ATP7A overlapping and juxtaposing with RAB11 demonstrating its presence in the RAB11 positive compartment. B (upper): ATP7A colocalizing with RAB11 positive compartments in the neurites. B (Lower): Magnified area from the upper panel shows colocalization and juxtaposition with RAB11.

RAB11 and TGN38+Golgin97 positive compartments overlap considerably in the perinuclear region of the differentiated glia (Fig 6 C,D, supplementary Fig 6). In addition to localizing on the overlapping compartments, ATP7A resides on exclusive RAB11 positive compartments in the perinuclear glia (Fig 6C,D). Such overlapping of TGN38+Golgin97 and RAB11 compartments is not observed in the neurites (Fig 7B, supplementary Fig 6) where ATP7A localizes only on RAB11 compartment (Fig 7). Together, the results demonstrate that ATP7A resides on both TGN38+Golgin97 and RAB11 positive compartments including the overlapping compartments in the perinuclear glia. With differentiation TGN38+Golgin97 and RAB11 positive compartment extend into neurites in the glia, where the compartments gain exclusivity (no overlap of TGN38+Golgin97 and RAB11). Among them, ATP7A localizes only on the RAB11 positive compartment in the neurites. Increased localization into the recycling endosomes, with differentiation, suggests role of these vesicles in either ready release or storage of the copper.

**Figure 6:**
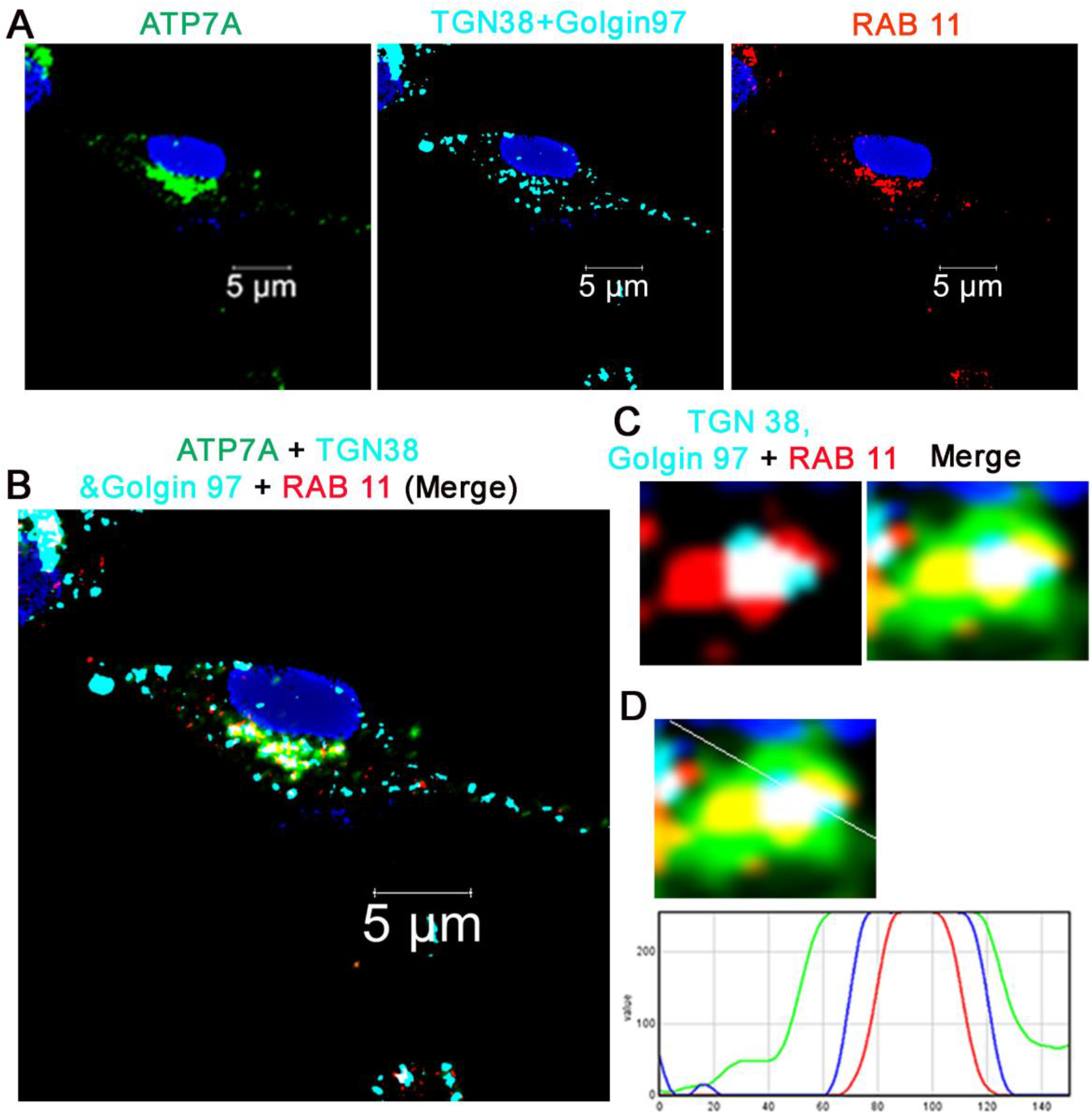
ATP7A resides both in the Trans Golgi Network (TGN38+Golgin97) and the recycling endosomes (RAB11) in the perinuclear(soma) region of the differentiated C-6 (glia) cells. A – Localization of ATP7A, TGN38+Golgin97 and RAB11 in the perinuclear region of the differentiated C-6 (glia). B – Merged (ATP7A, TGN38+Golgin97, RAB11) image demonstrating localization of ATP7A in the TGN38+Golgin97 and RAB11 positive compartments in the perinuclear region of the glia. C – An enlarged region showing exclusive and overlapping TGN38+Golgin97 and RAB11 positive compartments (left) and merged image (right) showing localization of ATP7A in the TGN38+Golgin97 and RAB11 positive compartments in the perinuclear region. D – Graph showing ATP7A distribution in the overlapping and exclusive RAB11 and TGN38+Golgin97 compartments in the perinuclear region.

**Figure 7:**
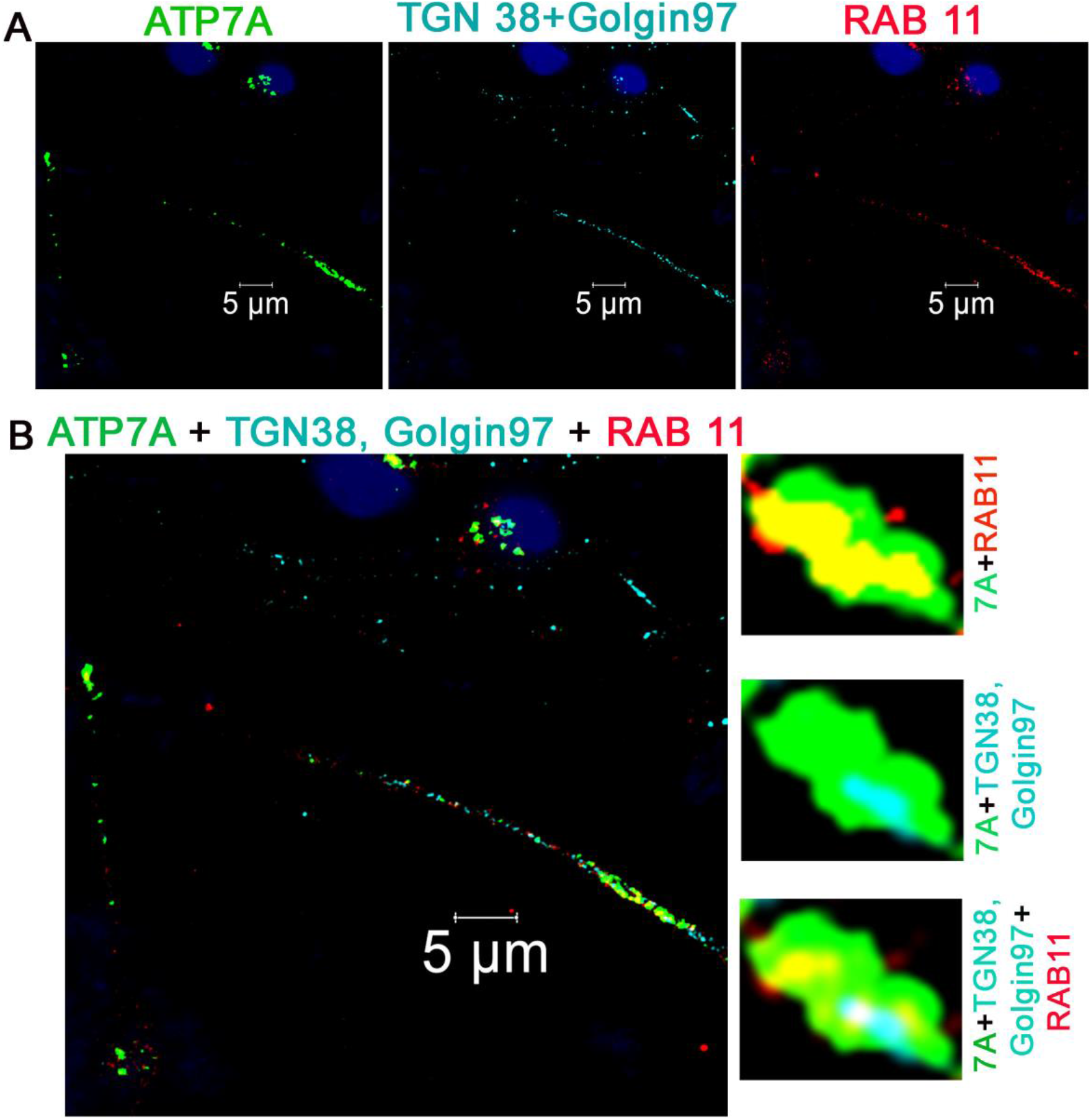
ATP7A colocalizes with only the recycling endosomes (RAB11) in the neurites of the differentiated C-6 (glia) cells. A – Localization of ATP7A, TGN38+Golgin97 and RAB11 in the neurites of the differentiated C-6 (glia). B (left) – Merged (ATP7A, TGN38+Golgin97, RAB11) image demonstrating colocalization of ATP7A (green) with RAB11 (red) and not TGN38+Golgin97 (cyan) in the neurite. The nuclei have been indicated in blue. B (right) – An enlarged region, in the neurite showing colocalization of ATP7A with RAB11 and lack of any colocalization of ATP7A with TGN38+Golgin97.

### (iv) Localization of the ATP7A into the RAB 11 positive vesicles in the differentiated C-6 (glia) cells is dependent on the level of intracellular copper

ATP7A localizes into the RAB 11 positive recycling endosomes in the differentiated glia. We wanted to investigate if this vesicular localization of ATP7A, along the glial processes, is copper dependent. To investigate this, C-6 cells were differentiated and treated for 2 hours with copper chelator ammounium tetra thiomolybdate (TTM) to chelate intracellular copper. Copper chelation led to the disappearance of the ATP7A on the vesicular structures and localized it entirely into the TGN (Fig 8). This suggests that the vesicular localization of ATP7A is dependent on the intracellular copper level. Further, we investigated whether this vesicular ATP7A is crucial for the glial differentiation. C-6 cells were plated, grown, and differentiated in the presence of TTM. Copper chelation did not affect the differentiation as is evident from the neurite generation (Fig 9A) and unaffected level of GFAP mRNA (Fig 9B). These cells also showed complete depletion of the vesicles in the glial processes with colocalization of ATP7A with TGN38+golgin97 (Fig 9C). This suggests that the localization of ATP7A into the RAB11 positive vesicles, in the glial neurites, do not play role towards the differentiation of the glial cells.

**Figure 8:**
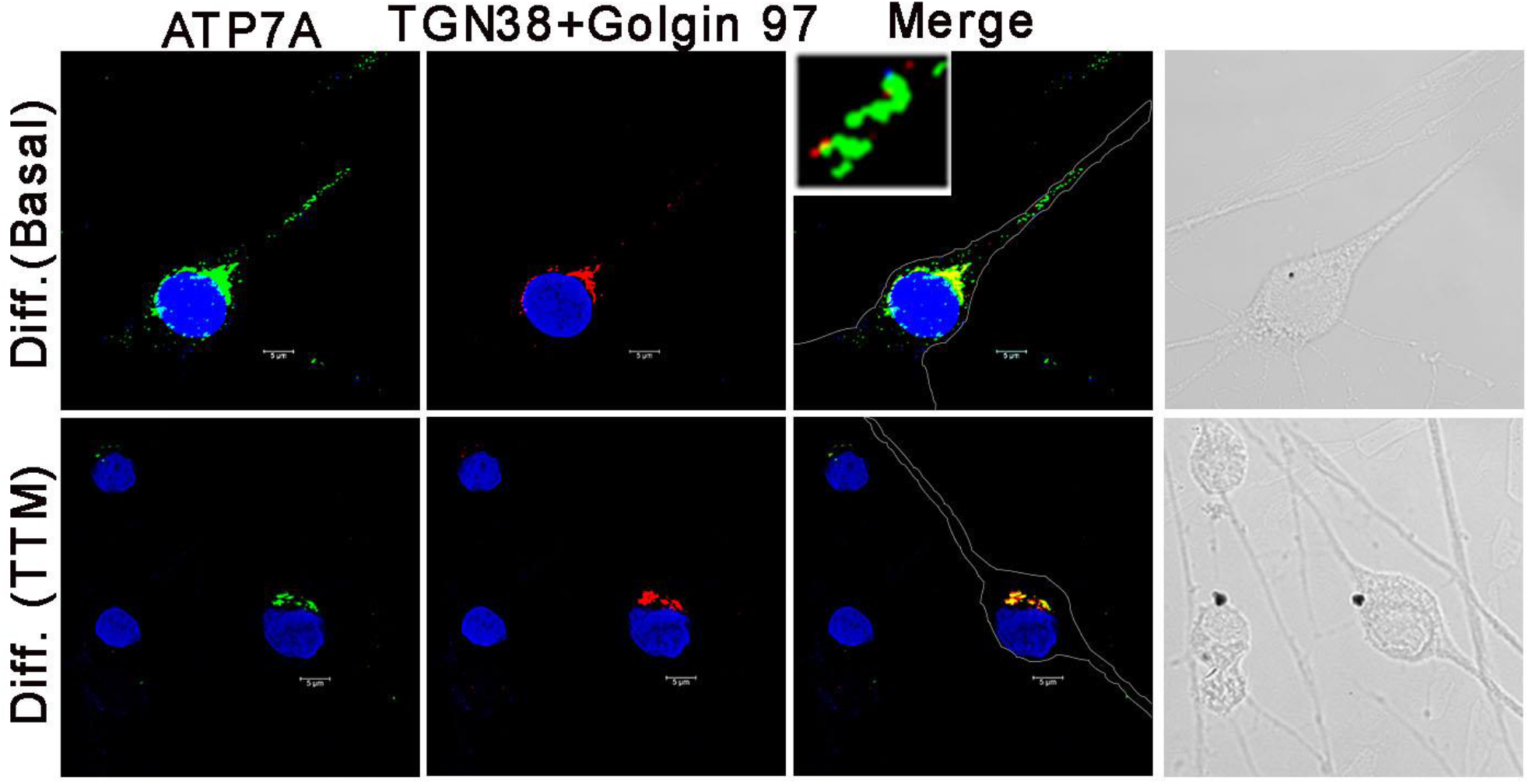
Presence of the ATP7A containing vesicles in the neurites of the differentiated C-6 (glia) cells is dependent on the intracellular copper level. C-6 cells were differentiated as described, and then treated with 30uM TTM for 2 hours. Disappearance of the ATP7A containing vesicles along the neurites and retention of ATP7A in the TGN38 and Golgin97 positive Trans Golgi Network (TGN) is observed in response to the copper chelation.

**Figure 9:**
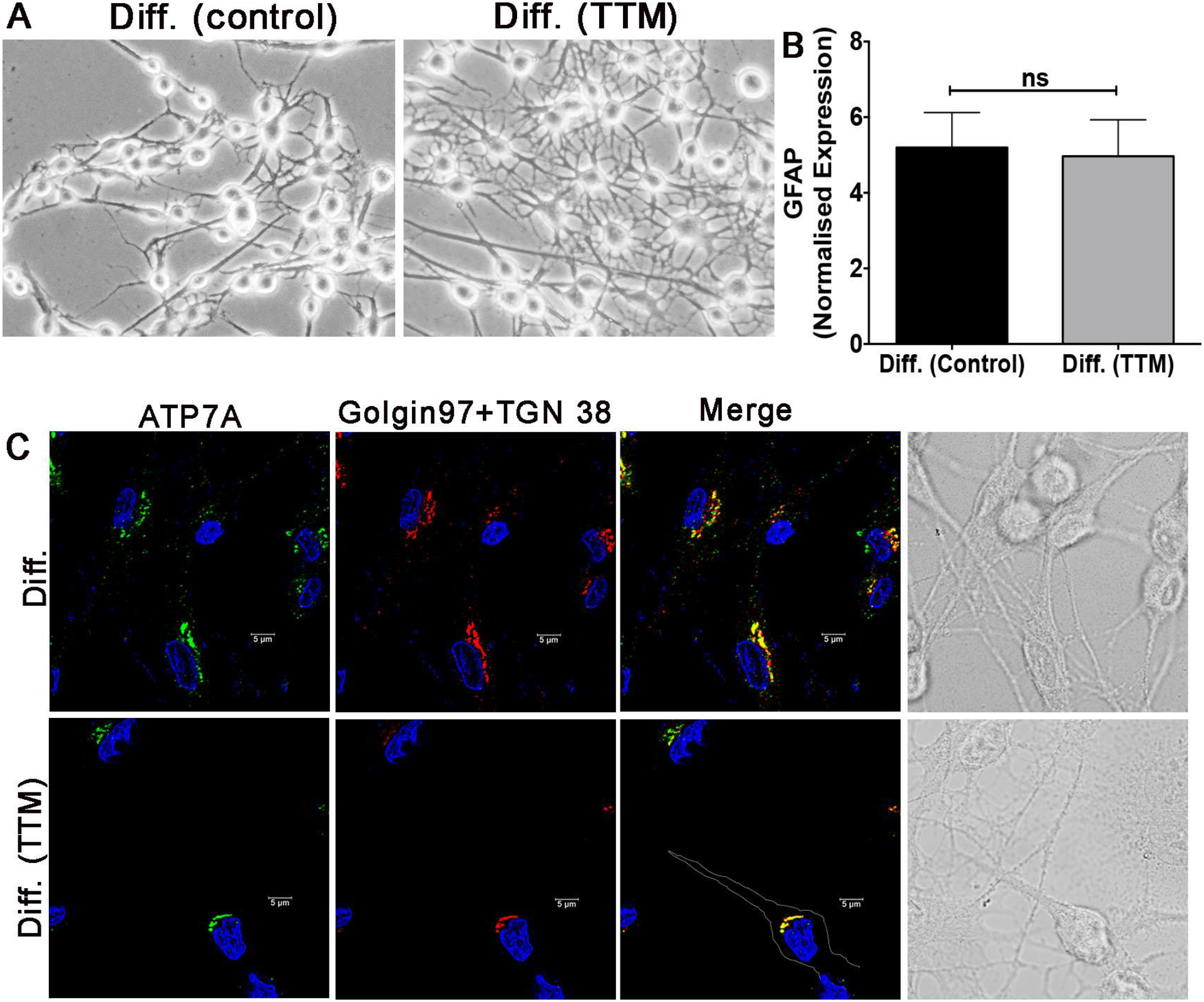
Chelation of the intracellular copper does not affect the differentiation of the C-6 cells into glia. C-6 cells are plated and differentiated in the presence of 30uM Ammonium Tetrathiomolybdate (TTM). A – Bright field image of the C-6 cells differentiated in presence and absence of TTM. B – Relative mRNA level of GFAP in the glial cells differentiated in presence and absence of TTM (no of replicate – 3). C - ATP7A localization in the glial cells differentiated in presence and absence of TTM.

### (v) Induction of the copper dependent ERK1/2 phosphorylation during the neuronal and not the glial differentiation

Retention of ATP7A into the TGN despite high intracellular copper suggests role of the copper towards the differentiation of the PC-12 cells into neurons. Activation of ERK1/2, through phosphorylation, has been observed during the differentiation of the embryonic stem cells into neurons (18). ERK1/2 activation is dependent on two Cu(I) ions binding to its upstream activator MEK1/2 (19). We wanted to investigate the status of the ERK1/2 phosphorylation during the differentiation of neuronal and glial cells. PC-12 derived neurons show an increased ERK1/2 phosphorylation (Fig 10A). We wanted to investigate if ERK1/2 phosphorylation accompanies neuronal differentiation and is not just associated with the differentiation of the PC-12 cells. To confirm this, we investigated ERK1/2 phosphorylation during the differentiation of the human fetal brain derived neuronal progenitor cells. These neurons also showed increased ERK1/2 phosphorylation like the PC-12 derived neurons (Fig 10B). This suggests that the increased intracellular copper, in the neurons is utilized towards activating pathways essential for the differentiation and ERK1/2 activation is one of them. Interestingly we observed 3 times downregulation of ERK1/2 phosphorylation in the glia compared to the undifferentiated C-6 cells (Fig 10C). Therefore, we observed opposite effects on ERK1/2 phosphorylation during the neuronal and glial differentiation. The experiments demonstrate that intracellular copper is not crucial for the glial differentiation. This is evident from the low intracellular copper in the differentiated glia and the glial differentiation not affected by the copper chelation. Therefore, the implication of the presence of copper dependent localization of ATP7A in the RAB11 positive vesicles might be either a ready release and/or rendering the copper unavailable for the pathways like ERK1/2 phosphorylation.

**Figure 10:**
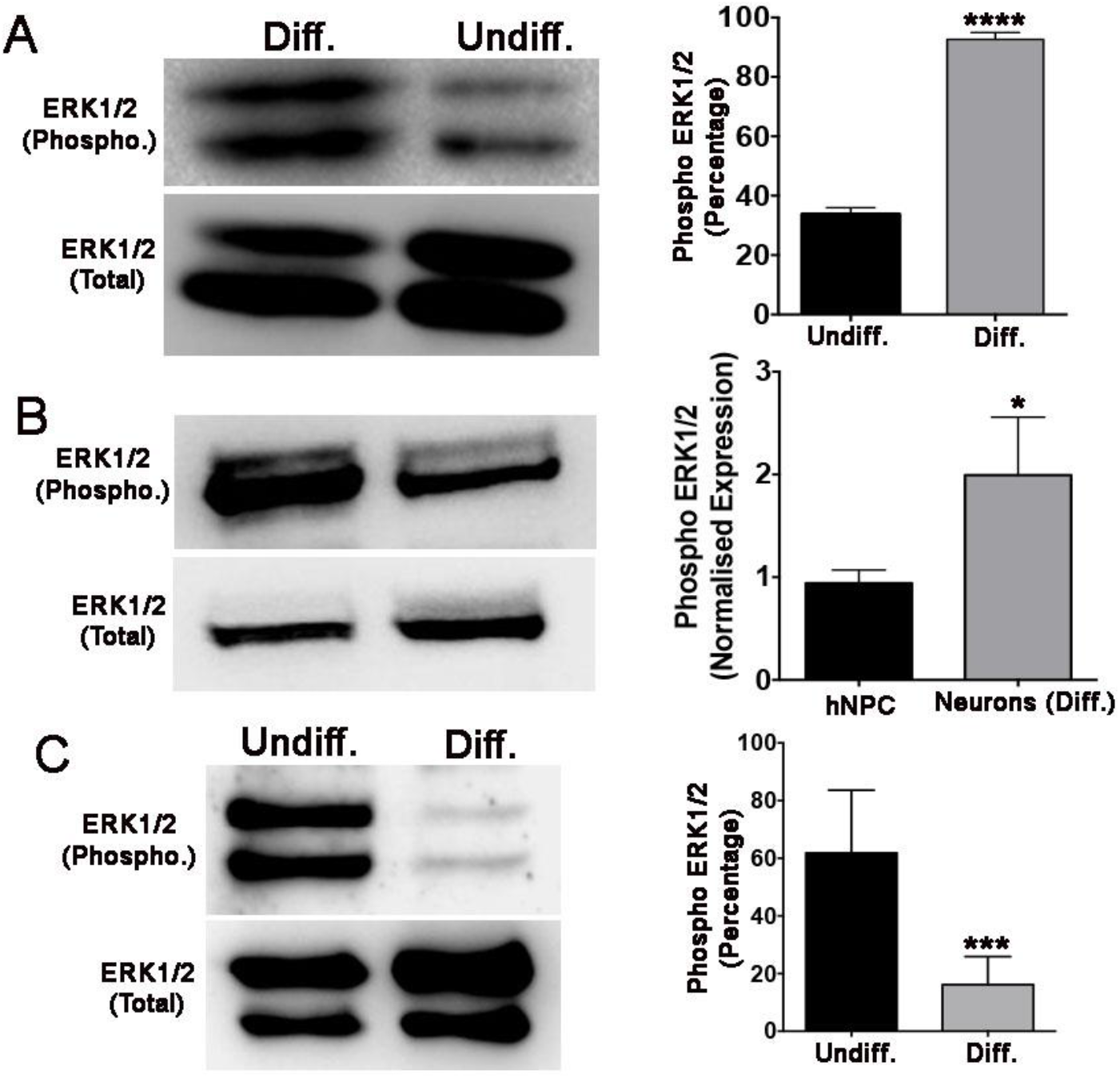
ERK1/2 phosphorylation in the differentiated neurons and glia. A – Increased ERK1/2 phosphorylation in the differentiated PC-12 (neuron) cells. B – Increased ERK1/2 phosphorylation in the neurons differentiated from the human fetal brain derived <u>N</u>euronal <u>P</u>rogenitor <u>C</u>ells (NPC). C – Decreased ERK1/2 phosphorylation in the differentiated C-6 (glia) cells.

### (vi) Intracellular copper chelation reduces ERK1/2 phosphorylation and neurite formation in the differentiated PC-12 (neuron) cells

Since the differentiation of neurons involves ERK1/2 phosphorylation (18), and ERK1/2 phosphorylation is copper dependent (20), we wanted to investigate if the increased intracellular copper in the differentiated PC-12 cells is utilized towards ERK1/2 phosphorylation. We investigated the effect of chelation of intracellular copper on ERK1/2 phosphorylation and differentiation of the PC-12 derived neurons. Chelation of the intracellular copper during the differentiation by treatment with TTM does not affect the differentiation as is evident from the unchanged neurite length (supplementary Fig 7A,B). Also, ERK1/2 phosphorylation does not decrease under such condition (supplementary Fig 7D). We hypothesized that the treatment with 10μM TTM at the onset of differentiation could not chelate the bioavailable intracellular copper. Therefore, we treated the PC-12 cells with higher dose (30μM) of TTM during the differentiation. But under this condition also there was no effect on the differentiation of the PC-12 cells (supplementary Fig 7A lower panel, right). Estimation of the intracellular copper level demonstrated that the treatment with 10uM TTM during differentiation could only reduce 29% of the intracellular copper (supplementary Fig 7C), in presence of the high expression of CTR1. This suggests that chelation of intracellular copper at the onset of differentiation is unable to chelate bioavailable Cu(I) and also remove Cu(I) from the MEK1/2 thereby failing to affect ERK1/2 phosphorylation.

Inorder to efficiently reduce copper mediated ERK1/2 activation, we cultured PC-12 cells for 48 hours in 10μM TTM and differentiated these cells in presence of copper chelator. Estimation of intracellular copper, under this condition, demonstrated decrease in 77% of the intracellular copper thereby suggesting depletion of the bioavailable Cu(I) (supplementary Fig 7C). Under this condition, we observed that copper chelation decreased neurite length at the onset of differentiation (T1- 48 hours after the first treatment with NGF) (Fig11 A,B). This data demonstrates that intracellular copper triggers pathways crucial for the generation of the neurites. Decreased expression of the B-III Tubulin (Fig 11C) in these cells confirms that the chelation of the intracellular copper affects the differentiation of the PC-12 cells. Neurite outgrowth is observed in the T1phase (Fig 11B) of the differentiation. However, we observed no change in the ERK1/2 phosphorylation in the T1 phase (Fig 11D, right). But we observed 4-fold (approximately) increase in the ERK1/2 phosphorylation in the T2 (Fig 11D, right). This suggests that ERK1/2 phosphorylation might not play role in the generation of the neurites. Chelation of intracellular copper led to two-fold decrease in the ERK1/2 phosphorylation in the T2 (Fig 11D) confirming that the intracellular copper, in the neurons, is also utilized towards phosphorylating ERK1/2. Therefore, the neuronal copper is not solely utilized towards the ERK1/2 phosphorylation. Our data indicates that in addition to ERK1/2 activation, intracellular copper triggers pathways crucial for the neurite outgrowth. Besides a bulk of the cytosolic copper might also be transported to the secretory pathway as demonstrated (9). Based on the observed increase in the ERK1/2 phosphorylation in the T2 phase of differentiation, we hypothesized its role towards the neuronal viability.

**Figure 11:**
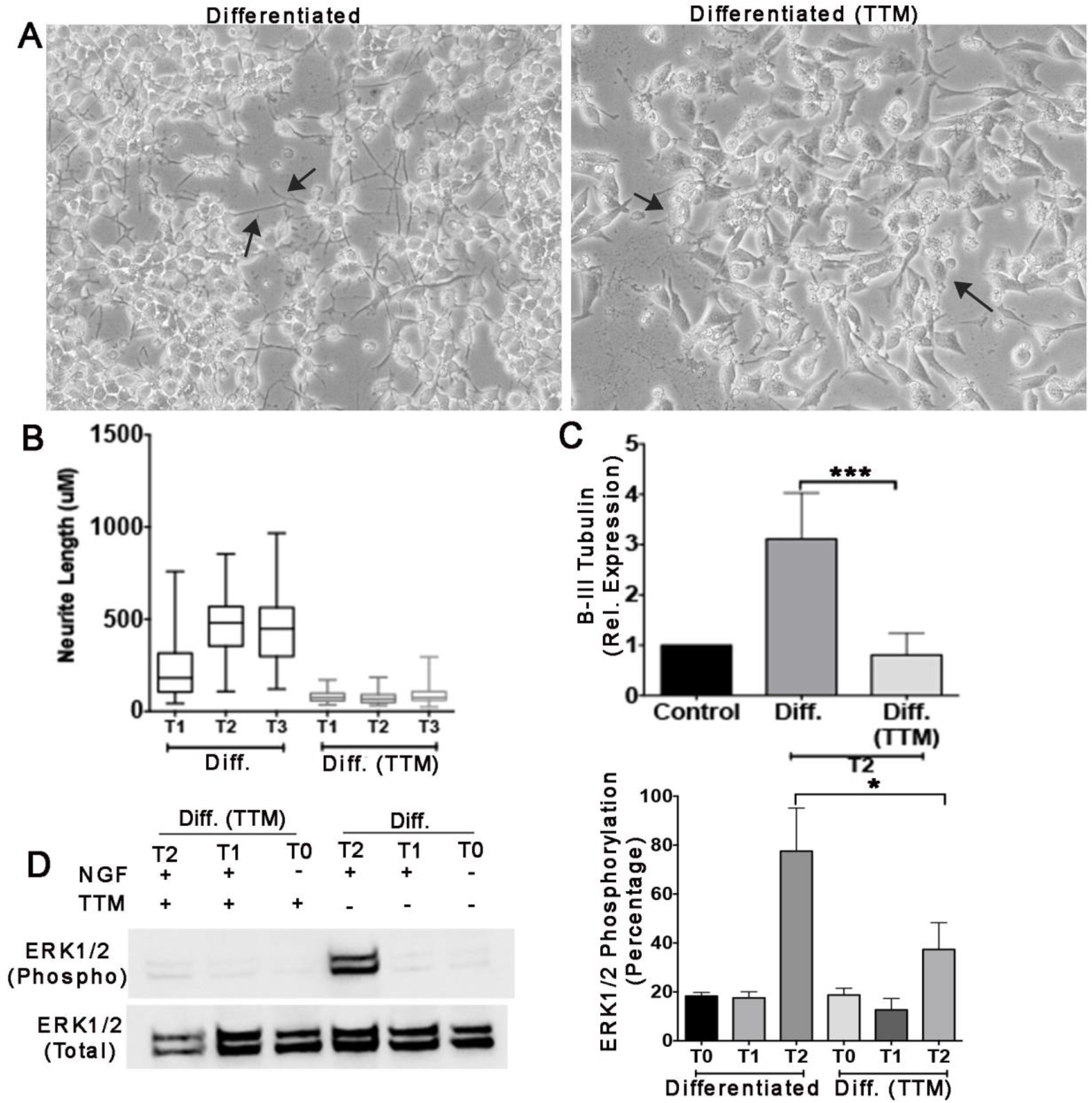
Copper chelation affects the neurite outgrowth and ERK1/2 phosphorylation in the differentiated PC-12 (neurons) cells. A - Bright field image of the cells cultured and differentiated in the presence and absence of TTM. B –Quantitative representation of the neurite length in the PC-12 cells cultured and differentiated in the presence and absence of TTM. C – Relative mRNA for B-III tubulin in the undifferentiated and cells differentiated under similar conditions as above. D – Western immunoblot (left) and quantitation by densitometry (right) demonstrating ERK1/2 phosphorylation in the PC-12 cells differentiated under the same condition (number of replicates – 3). T0, T1, T2 denote the different time points of the NGF treatment. T0 – no treatment, T1 –first treatment with NGF, T2 – second treatment with NGF at an interval of 48 hours after the first one.

### (vii) U0126 mediated inhibition of the ERK1/2 phosphorylation decreases the viability of the PC-12 derived neurons

ERK1/2 phosphorylation is significantly activated in theT2 phase of differentiation of the PC-12 cells (Fig 11D). Since ERK1/2 is not activated at the initial phase along with the neurite generation, we hypothesized that it plays role in the neuronal survival rather than neurite generation. We investigated effect of the inhibition of ERK1/2 phosphorylation by the MEK1/2 inhibitor, U0126 on the PC-12 derived neurons. PC-12 cells (undifferentiated) were treated with 2-8μM MEK1/2 inhibitor U0126. The treatment did not affect the viability of the undifferentiated cells (supplementary Fig 8A). A significant concentration dependent inhibition of the ERK1/2 phosphorylation was observed in the PC-12 derived neurons when differentiated in the presence of U0126 (supplementary Fig 8B). These neurons showed significantly decreased viability (Fig 12 A,B). Quantitation of the viable cells, estimated by the neutral red incorporation assay, demonstrated 57-84% decrease in the viability of the neurons (Fig 12B) during the T2 phase, in response to the U0126 treatment. Decrease in the B-III tubulin expression (Fig 12C) confirms the selective loss of the neuronal cells in response to the treatment.

**Figure 12:**
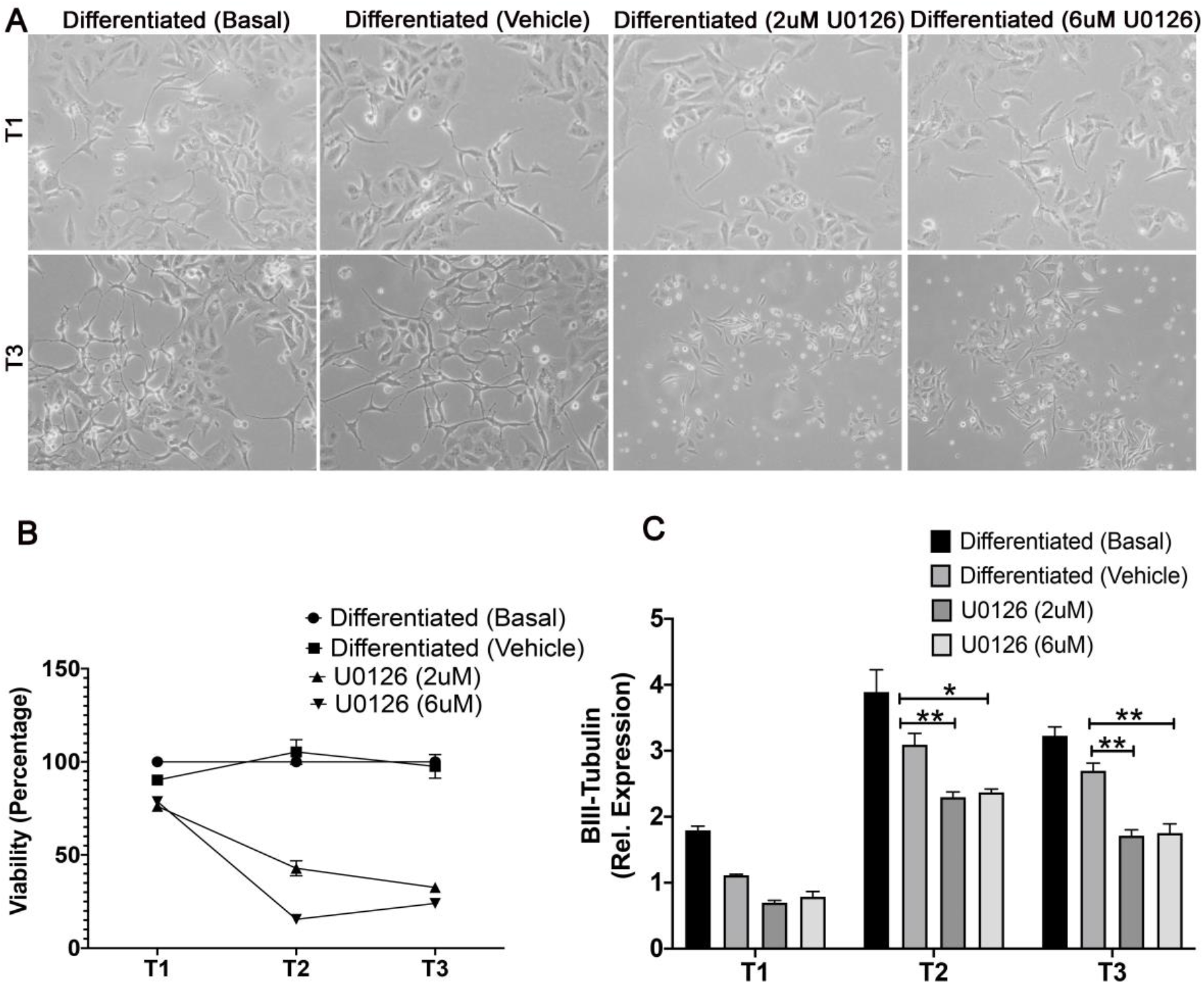
Inhibition of the ERK1/2 phosphorylation by U0126 affects the viability of the differentiated PC-12 (neurons) cells. A –Bright field images of the PC-12 cells differentiated in the presence or absence of the U0126. B – Percentage viability of the PC-12 cells differentiated in the presence or absence of the U0126. C – Relative mRNA expression of B-III tubulin in the PC-12 derived neurons under similar conditions as mentioned.

## Discussion

This is the first comparative study of the changes in the cellular copper homeostasis during the neuronal and the glial differentiation. Although the existing cell-based model might not exactly mimic the *in vivo* situation, yet they provide a glimpse of the role of cellular copper homeostasis during the neuronal and glial differentiation. We have observed the differential role of intracellular copper in triggering the pathways in the neurons and the glia during differentiation. Differentiation of the PC-12 cells involve increased intracellular copper, whereas, differentiated glia have low intracellular copper (Fig 13). Requirement of copper for the differentiating neurons is evident from the increased level of CTR1 mRNA accompanying the differentiation and decreased neurite generation in response to copper chelation. However, ATP7A transcript remains unchanged with the differentiation of the PC-12 cells into neurons. The expression of ATP7A has been observed to increase with the differentiation of the SH-SY5Y cells (9). Interestingly ATP7A is retained into the TGN despite high intracellular copper in the PC-12 derived neurons (Fig 13). This localization suggests utilization of the high intracellular copper towards the differentiation. The intracellular copper has multiple utilizations. The present study demonstrates that the copper is utilized towards triggering pathways crucial for the neurite generation and ERK1/2 activation (Fig 13). The cytosolic copper, in the neurons, is also transported to the Golgi based secretory pathway (9). A part of the intracellular bioavailable copper might also be transported to the mitochondria (present study) based on the increased mRNA for COX17 in the PC-12 derived neurons.

**Fig 13:**
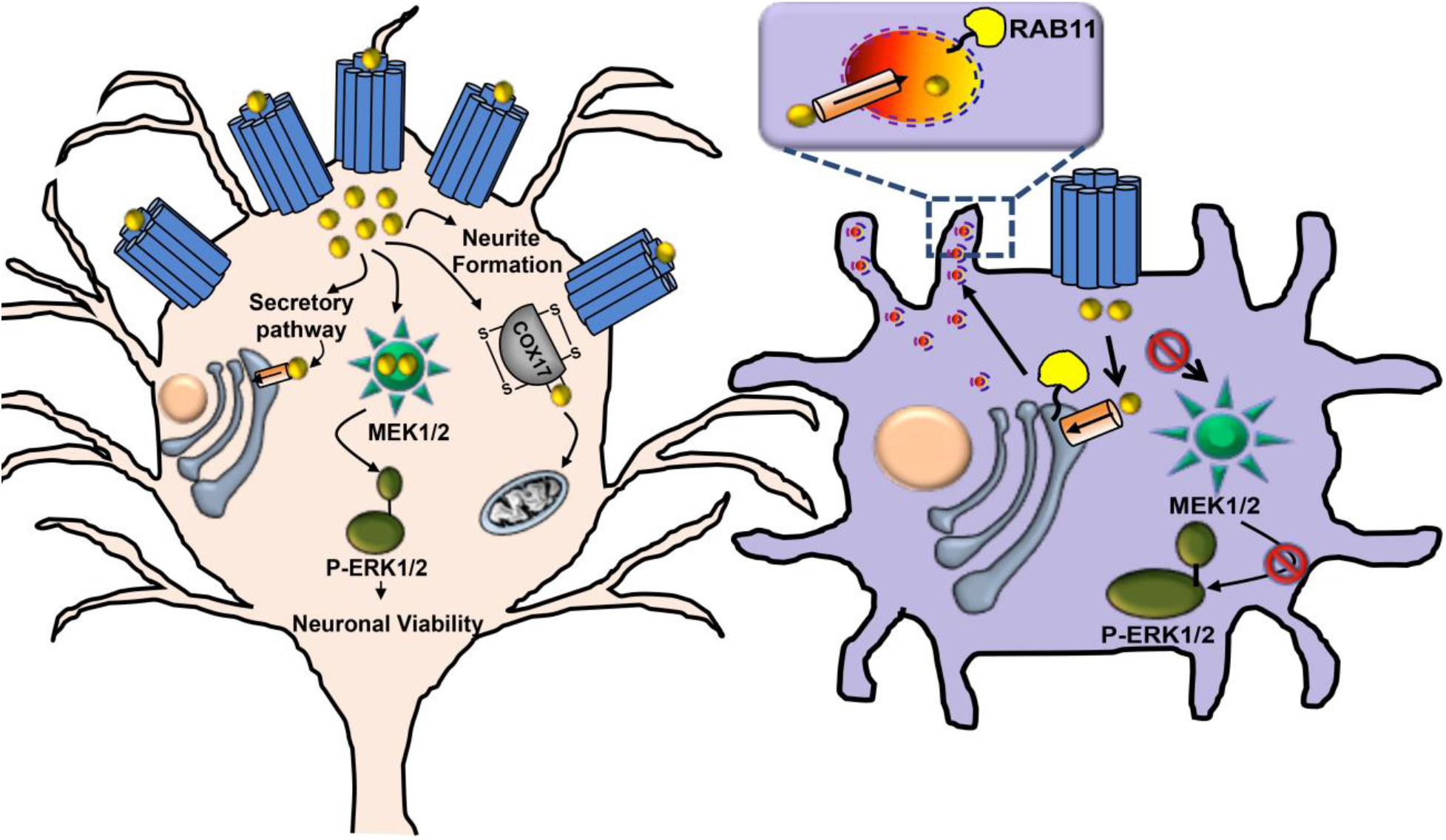
Utilization of the intracellular copper during the PC-12 (neuronal) and C-6 (glial) differentiation. Left: Differentiation of the PC-12 cells involve increased level of the CTR1 (blue cylinder) resulting into an increased intracellular copper (golden sphere). A bulk of the cytosolic copper is reportedly utilized by the Golgi based secretory pathway. Cytosolic copper binds to MEK1/2 (green circle with spikes) and triggers ERK1/2 phosphorylation (green ellipse) which is crucial for the viability of the neurons. ATP7A (orange cylinder) is retained in the TGN, despite high intracellular copper. Right: Differentiated C-6 (glia) have low intracellular copper and thereby downregulation of the copper dependent ERK1/2 phosphorylation. ATP7A localizes in the TGN38 and RAB11 positive compartments in the perinuclear region. In addition, ATP7A localizes into the RAB11 positive vesicles in the neurites (inset) despite low intracellular copper.

We report, for the first time, copper dependent ERK1/2 activation during the differentiation of the PC-12 and human fetal brain derived neuronal progenitor cells. Therefore, ERK1/2 activation is one of the cytosolic pathways triggered by the intracellular copper during the differentiation of the neurons. Copper dependence of ERK1/2 phosphorylation has already been demonstrated in melanoma cells (19,20) and further confirmed by our experiments in the neurons. Inhibition of the ERK1/2 activation by U0126 affecting the viability of the neurons highlights the importance of this copper dependent pathway towards the neuronal differentiation. ERK1/2 phosphorylation has been observed to be associated with differentiation of the Embryonic Stem (ES) cells into neurons (18). But the present study demonstrates the role of the process towards the neuronal differentiation. Moreover, we demonstrate opposite status of the copper level and ERK1/2 phosphorylation during the neuronal and glial differentiation (Fig 13).

The glia, having low intracellular copper, have ATP7A localizing into both TGN (perinuclear) and RAB11 (neurites) positive vesicles (Fig 13). The vesicular localization of ATP7A into the RAB 11 positive compartments is reported for the first time. Presence of these vesicles at low intracellular copper, is intriguing. ATP7A expression has been reported in the developing axons in the olfactory neurons (21). Extensions of the Golgi compartments and TGN38 has been detected in the endosomes (10) in the neurons. Mannose-6-phosphate receptor recycles between TGN and late endosomes in the neuronal processes (22). But ATP7A containing vesicles, identified in the glia, do not overlap with any of the classical TGN markers. Also RAB11 positive recycling endosomes have been implicated in neurite extension in the PC-12 derived neurons, dorsal root ganglia and Hippocampal neurons (23). But we did not observe such vesicles in the neurons. It is observed only in the C-6 derived glia. Chelation of the intracellular copper in the glia, leading to the disappearance of these compartments, does not affect the glial differentiation. Therefore, the primary function of these vesicles might be to store copper either for supply to the neurons or to bypass the cytosolic copper thereby making it unavailable for the MAP kinases (Fig 13). Keeping in mind the already implicated role of the glia in maintaining ion homeostasis (24), we hypothesize that the glia might be the key regulator of copper homeostasis in the brain. The ability of the glial cells to sequester copper into RAB11 positive vesicles underlies its ability to reportedly (25) withstand copper induced toxicity. Role of cellular copper homeostasis has earlier been suggested to play role in neuronal differentiation. However, intracellular copper has been hypothesized to be diverted towards the secretory pathway aiding into the differentiation of the SH-SY5Y cells (9). But the role of the intracellular copper to be utilized only towards the secretory pathway via TGN during differentiation might not be the only pathway of copper utilization during the differentiation. Our study demonstrates a direct route of copper utilization during neuronal differentiation, in addition to incorporation into the cuproenzymes.

Overall, we demonstrate that copper plays important role in both neuronal and glial differentiation.

### Experimental Procedures

#### Cell Culture, differentiation

The PC12 cells are differentiated as reported (26) with minor modifications. Cells maintained in in DMEM (DMEM, high glucose, GIBCO) containing 10% horse serum (heat inactivated, GIBCO), 5% Fetal Bovine Serum (Heat Inactivated, HiMedia), 1% Pen-Strep (HiMedia), 1% Amphotericin-B (GIBCO) at 37^0^C and 5% CO2. The C-6 glioma cells are maintained in DMEM containing 10% FBS, 1% Pen-Strep,1% Amphotericin-B at 37^0^C and 5% CO2. For differentiation of PC-12 cells 2.5×10^5^ are plated on each well of a 6-well plate coated with collagen and cultured overnight. The cells are rinsed twice with phosphate-buffered saline. Differentiation is initiated by adding 100ng/ml Nerve Growth Factor (Sigma Aldrich) in PC12 priming media containing DMEM and 0.5%-1% horse serum. Three subsequent treatment are performed, first two at an interval of 48h and the last one at 24h after the second treatment. C-6 cells are differentiated as mentioned (27) with minor modifications. For differentiation of glia, C6 cells (1X10^5^ cells) are plated on each well of a 6-well plate and cultured for 48h (approximately) to 50-60% confluency. The cells are then rinsed with phosphate-buffered saline and cultured in DMEM medium without fetal bovine serum for 1 hour. Differentiation is initiated by adding 500μM dBcAMP (N^6^,2′-O-Dibutyryladenosine 3′,5′-cyclic monophosphate sodium salt, Sigma Aldrich) and 1mM Theophylline (MP Biomedicals) in culture medium without serum. Bright field images of undifferentiated and differentiated cells are acquired using DM IL LED Fluo (Leica microsystems) and Carl Zeiss^™^ PrimoVert^™^ Inverted Microscope (ZEISS).

#### Human fetal neural stem cell culture and differentiation

Human fetal brain tissue is isolated from 10-15 week old aborted fetuses according to guidelines of Institutional Human Ethics and Stem Cell and Research Committee of National Brain Research Centre in compliance with the approvals of Indian Council of Medical Research. The aborted tissue is used for research purpose only after obtaining the informed consent of the mother. Fetuses selected for the culture were healthy and had no sign of aneuploidy. The tissue is processed for neural stem cell culture as described previously (28), (29). Briefly, NSCs are isolated from the telencephalon of the fetal brain and are cultured onto the Poly-D-Lysine (Sigma-Aldrich) coated culture-wares in neurobasal medium (Invitrogen). Media is added with following: Neural Survival Factor-1 (Lonza), N2 supplement (Invitrogen), Bovine Serum Albumin (BSA) (Sigma-Aldrich), L-glutamine (Sigma), fibroblast growth factor (25 ng/ml) (bFGF) (Peprotech), epidermal growth factor (20 ng/ml) (EGF) (Peprotech), penicillin and streptomycin solution (Invitrogen) and gentamycin (Sigma).

Neural stem cells (NSCs) are passaged at least nine times before using them for experiments. NSCs were characterized for the expression of stemness markers- SOX2 and NESTIN by immunocytochemistry and PCR. More than 99% of the cells showed immunoreactivity towards stemness markers- NESTIN and SOX2. Also, cells are analysed for lineage-specific markers-GFAP (astrocytic marker) and MAP2 (neuronal marker), more than 95 % cells are negative for GFAP and MAP2 when maintained in above-mentioned stem cell medium. For differentiation of NSCs into neurons, EGF and bFGF are replaced with brain-derived neurotrophic factor (10ng/ml) (BDNF) (Peprotech) and platelet-derived growth factor (10ng/ml) (PDGF-AB) (Peprotech) for 21days. Post differentiation cells are assessed for expression of MAP2 and Tuj-1; more than 95% cells are positive for these neural markers.

##### Measurement of intracellular copper by Inductively Coupled Plasma-Mass Spectrometry (ICP-MS) and ICP-OES (Indictively Coupled Plasma – Optical Emission Spectrometry)

PC-12 Cell (undifferentiated and differentiated) pellets are washed with ice cold PBS and solubilized in RIPA Lysis buffer (10mM Tris-Cl pH 8.0, 1mM EDTA, 0.5mM EGTA, 140mM NaCl, 1% Triton X-100, 0.1% sodium deoxycholate and 0.1% Sodium dodecyl sulphate and protease inhibitor cocktail) and centrifuged at 600Xg in 4^0^C to remove the cell debri and unlysed cells. Lysate is digested in 50% extrapure nitric acid (Finar) at 55°C for an hour. Digested sample is diluted with water to a final concentration of 3% nitric acid. Metal estimation analysis is performed using an Thermo Fisher Scientific X-Series 2 ICP-MS. Data were quantified using a SRM2711a – Montana II soil moderately elevated trace element concentration standard (NIST – National Institute of Standards and Technologies). Copper concentration is normalized to total protein and represented.

For estimation of the total copper in the undifferentiated and differentiated PC12, in response to the TTM treatment, samples are similarly prepared as mentioned. The digested samples are filtered through 0.45 micron syringe filter. Copper is estimated by iCAP 7000 Series ICP-OES (Thermo Scientific). Copper concentration is normalized to the total protein and represented. Each result was replicated at least for 3 times.

Undifferentiated and differentiated C6 cells are washed twice with Phosphate Buffered Saline (pH:7.5), trypisinized, centrifuged and collected as pellets. The pellets are digested with 2% suprapur HNO3 (Merck) at a concentration of 3ml /0.25 mgs of the cell pellet for an hour at room temperature. The cell pellets are further digested at 100^0^C for an hour in a microwave (Microwave digestion condition: Power=800 W; Temperature=100^0^C; Hold time= 1 h). The digested sample is filtered through 0.45 micron syringe filter. Elemental copper standard is prepared by digesting copper foil (Alfa Aeasar) with suprapur HNO3 under similar condition. Metal estimation has been done using Inductively coupled Plasma - Optical Emission Spectrometry (ICP-OES) using iCAP DUO 6500 (Thermo Scientific). Copper concentration is normalized to total protein and represented. Each result was replicated at least for 5 times.

##### Measurement of mRNA level by Real Time PCR

Undifferentiated and differentiated cells are cultured in 60 mm dish. Cells are harvested and washed with phospate buffered saline. RNA was extracted with the Nucleospin RNA isolation kit (Macherey-Nagel) according to the manufacturer’s instruction manual. The cDNA is prepared by reverse transcription using iScript^™^ Select cDNA Synthesis Kit (Biorad Laboratories and TaKaRa) using 1μg of extracted mRNA in a final volume of 20 μl by using anchored oligo(dT) and random hexamer primer following manufacturer’s instructions with minor modifications. Real Time PCR is performed using 2μl of prepared cDNA in a 10μl reaction volume with iTaqUniverSYBR Green SMX 1000 (Biorad Laboratories) using Bio-rad CFX96 touch Real Time PCR instrument. The primers are used to amplify the respective rat cDNA (supplementary table1). RPL29 cDNA is used as the reference gene.For PC-12 cells, the PCR included an initial denaturation at 95°C for 1 minute followed by denaturation at 95°C for 30 seconds, annealing and extension at 60°C for 60 seconds (32 cycles). For C-6 cells, initial denuration is carried out for 30 seconds followed by denaturation for 5 seconds and annealing and extension for 30 seconds (40 cycle). Relative mRNA abundance is estimated by 2^-ΔΔCT^method. Each experiment is repeated at least thrice.

##### Immunofluorescence and image acquisition to investigate localization of ATP7A in the neurons (PC-12) and glia (C-6)

Undifferentiated and differentiated cells are cultured directly (C-6) or on collagen (PC-12) coated coverslips. Cells are first washed twice in 1X phosphate buffered saline. Cells are fixed with 4% paraformaldehyde in phoshate buffer saline for 3 minutes on ice followed by chilled methanol and blocked with 3% (w/v) BSA. Coverslips with cells are inclubated in primary antibodies in 1% Bovine Serum Albumin (Albumin Bovine (pH 6-7) fraction V, 98%, Sicco Research Laboratories Pvt. Ltd.) overnight. Rabbit polyclonal Anti-ATP7A antibody (abcam, 1:300 dilution), mouse monoclonal TGN38 antibody (Novus Biologicals, 1:400 dilution), mouse monoclonal Golgin 97antibody (Thermo Fisher Scientific, 1:400 dilution), mouse monoclonal LAMP1 (Developmental Studies Hybridoma Bank, University of Iowa1:400 dilution,), mouse monoclonal Gama Tubulin (Invitrogen, 1:400 dilution), mouse monoclonal RAB 7 (Novus Biologicals, 1:50 dilution), goat polyclonal RAB11 (Santa Cruz Biotechnology Inc., 1:50 dilution), rabbit monoclonal anti-FLAG antibody (Cell signalling technology, 1:600), mouse monoclonal anti-GFAP (Sigma Aldrich) is used to detect the respective proteins. Goat anti-rabbit Alexa Flour 488 (abcam), donkey anti-mouse Alexa Flour 568 (ThermoFisher), donkey anti-goat Alexa Flour 568 (ThermoFisher), donkey anti-mouse Alexa Flour 633 (Thermo Fisher). Secondary antibodies are used at 1:2000 dilution for detection. Cell are mounted with Fluoroshield mounting medium (Sigma) (Containing DAPI) and imaged with a confocal microscope (SP8 Lightening confocal microscope, Leica Microsystems). Image is acquired using oil immersion 63X objective and deconvoluted using Leica application suiteX Lightning software (Leica microsystems). Fluorescence intensity line profile plot is done using Image-J (MBF).

For Stimulated Transmission Emission Depletion microscopy, cells are first washed twice in 1X PBS for 3 minutes. Cells are fixed with 2% paraformaldehyde in PBS for 20 minutes at room temperature followed by the treatment with 50mM ammonium chloride for 20 minutes. Cells are blocked with 1% (w/v) BSA permiabilized with 0.075% saponin (Sigma Aldrich) in phosphate buffer. Cells are incubated in primary antibodies in 0.5% overnight at 4 degrees. Cells are incubated in secondary antibodies for 90 minutes. Donkey anti-mouse Alexa Fluor 633 at 1:1000 dilution, is used as secondary antibody for detection of RAB-7. Rest of the secondary antibodies (1:1000) used is like those for confocal microscopy. Cell are mounted in Prolong (R) goldmounting medium (without DAPI) (Cell Signalling Technologies) and imaged with a confocal microscope (SP8 Lightning confocal microscope with STED attachment, Leica Microsystems).

##### Chelation of intracellular copper in the Glia (C-6) by treatment with TTM

For investigating if ATP7A vesicles in glia (C-6) contain copper, undifferentiated and differentiated cells are plated on coverslips and cultured as described. Cells are treated with 30μM Ammouniumtetrathiomolybdate (Sigma) for 2 hours. Cells were fixed, permeabilized, blocked and prepared for immunostaining as described in the previous section.

For investigating the effects of copper chelation on differentiation, glia (C-6) cells are plated on medium containing 30μM TTM and cultured for 36 hours. Differentiation is initiated (as described) in presence or absence of 30μM TTM. Cells are harvested and prepared for immunocytochemistry as described in the previous section.

For investigating effect of copper chelation on ERK1/2 phosphorylation and PC-12 differentiation, cells were plated on 6 well plates. Differentiation is performed (as described) in presence of 10uM TTM. Alternatively, PC-12 cells are plated on 6 well plates and cultured for 30 hours (30-40% confluency) in presence of 10μM TTM. Differentiation was performed (as described) in presence or absence of 10μM TTM. Cell are harvested 48hours after the second treatment with NGF.

##### Cell lysis and western immunoblot blot for detectection of Total and Phospho-ERK1/2 in Neuornal (PC-12), Glial (C-6 cells) and human fetal brain derived neurons

Undifferentiated and differentiated cells are cultured on 6 well plates. Cell pellets are resuspended with 100μl of RIPA lysis buffer and incubated on ice for one hour. Cells are homogenized with Dounce homogenizer with 220 strokes of the tight pestle. The homogenate is centrifuged at 600Xg for 10 minutes and supernatant is collected. Total protein in the lysate is estimated by Bradford assay. 60 μgms (PC-12 and C-6) and 40-50μgms (human fetal brain derived NPC and neurons) of lysate is mixed with gel loading dye (2% SDS, 2.5% ß mercapto-ethanol, 7.5% glycerol, 2M Urea and 0.005% bromophennol blue), boiled for 10 minutes at 95^0^C, kept at room temperature and resolved in 12% denaturing polyacrylamide gel. Protein is transferred to nitrocellulose membrane (BioRad) at 180mA current for 90 minutes. Membrane is blocked with 3% BSA at room temperature for 2 hours and incubated overnight at 4^0^C with primary antibody in 1% BSA and 0.1% tween-20. p44/42 MAPK (Erk1/2) (Cell Signalling Technologies) and Phospho-p44/42 MAPK (Thr202/Tyr204) Cell Signalling Technologies) (dilution of 1:3300) are used as primary antibody for detecting total and phopho-ERK1/2 respectively. Membrane is washed thrice with Tris buffer-saline (pH7.5) (TBS) and thrice with Tris buffer-saline containing 0.1% Tween-20 (TBS-T). Membrane is incubated in goat anti rabbit HRP conjucated secondary antibody (Sigma and Santa Cruz Biotechnology) for 2hours at room temperature. Membrane is washed thrice in TBS-T and thrice in TBS and chemiluminescence (clarity max, Biorad) detected using ChemiDoc^™^ Imaging System (170-01401) (Bio-Rad) instrument.

##### Neurite length estimation

In order to measure the length of the neurites of PC12 cells (undifferentiated and differentiated), bright field images of the cells are acquired and subsequently analyzed with the help of Image-J. In order to perform the analysis, resolution property (both in pixels and cm) of the image is acquired. The resolution of width/height of the image is obtained in pixel and centimeter. The size of each pixel in centimeter is obtained using this information. The image is then opened in ImageJ. The scale of the software is calibrated beforehand in accordance with the previously mentioned, calculation. This is done by providing the pixel to centimeter value in the ‘set scale’ menu bar under ‘Analyze’ option in Image-J. Input of pixel aspect ratio property of the image is also required. Following this, the length of the neurite lengths is measured with the help of the “line selection tool” in Image-J. The process of length measurement is repeated for at least in 50 cells. The average value is then downscaled to its appropriate and accurate unit (μM) by adjusting the magnifications.

##### Statistical analysis

Data were expressed as mean values +/- standred deviation. The Students paired t test is used to compare differences between two groups by GraphPad Prism 6 software (GraphPad Prism Inc, San Dieago,CA).statistically significant differences between them are indicated following the norms: *P<0.05,**P<0.01,and ***P<0.001.

##### Inhibition of the ERK1/2 phosphorylation by U0126 in PC-12 cells

For estimating the toxicity of U0126 on the PC-12 cells, 1.5×10^3 undifferentiated cells are plated on each well of 96 well plate. Cells are treated with varying concentrations (0-8 μM) of U0126 for 48 hours in complete medium.

For investigating the effect of ERK1/2 phosphorylation on PC12 differentiation, 5×10^4 cells are plated on 6cm dish and kept overnight. Cells are treated with or without U0126 (Cell Signalling Technology) in PC12 priming media (DMEM containing only 0.5% of horse serum) for two hours. After two hours, differentiation is initiated by 100ngm/ml of NGF in presence or absence U0126. Cells are differentiated as mentioned earlier in presence or absence of U0126. ERK1/2 phosphorylation is investigated by western immunoblot as described before.

For estimation of neuronal viability, PC-12 cells are plated at 1.25*10^4 cells in each well of a 12 well plate and grown for 18 hours. Cells are differentiated in the presence or absence of U0126.

#### Estimation of cell viability by Neutral Red Dye incorporation assay

PC-12 cells are cultured in presence or absence of U0126 as mentioned. Culture medium is aspirated as specific time points and cells are washed twice with Phosphate Buffered Saline (PBS) (pH:7.4) and added with 40ugm/ml Neutral Red in complete medium (DMEM containing 10% horse serum and 5% FBS) and incubated for two hours at 37^0^C and 5% CO2. Cells are gently washed with PBS followed by the addition of distaining solution (50% Ethanol and 1% Acetic acid) and absorbance recorded immediately at 540 nm either in a microplate reader (Varioskan LUX, Thermo Fisher) or a spectrophotometer (Eppendorf Biospectrophotometer, basic). Cell viability is expressed as percentage to the viability of the untreated differentiated cells.

## Funding and additional information

The study was supported by Ramanujan Fellowship Project (SB/S2/RJN-106/2015) and Extramural Research Project (EMR/2016/003293) to AB and Early Career Research Award (ECR/2015/000220) to AG from Science and Engineering Research Board, Department of Science and Technology, Government of India. The work is further supported by Wellcome Trust India Alliance Fellowship (IA/I/16/1/502369) and IISER- K intramural funding to AG.

## Author contributions

AB, AG and KC designed experiments. AB, KC, BR, AG, RB and PS wrote the manuscript. KC, SK, BR, RB, SD performed experiments. KC, SK, BR, AG participated in data acquisition, KC, AB and AG analyzed the data.

## Figures and Legends

## Supporting Information

**Supplementary figure 1:**
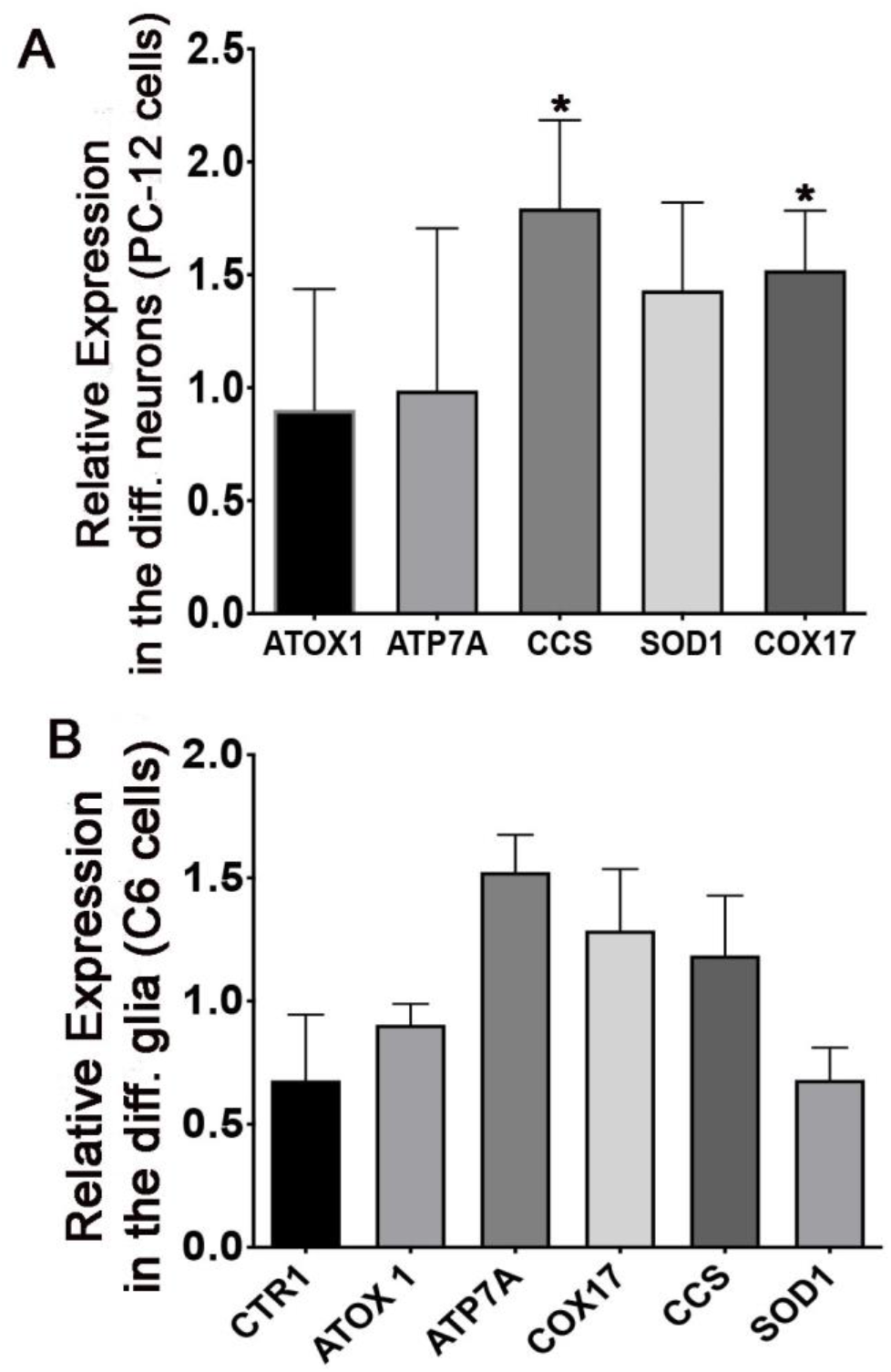
Relative mRNA abundance of the copper chaperones, transporters and cuproenzyme in the differentiated PC-12 (neurons) and C-6 (glia) cells. A: differentiated PC-12 (neurons) cells. B – Differentiated C6 (glia) cells.

**Supplementary figure 2:**
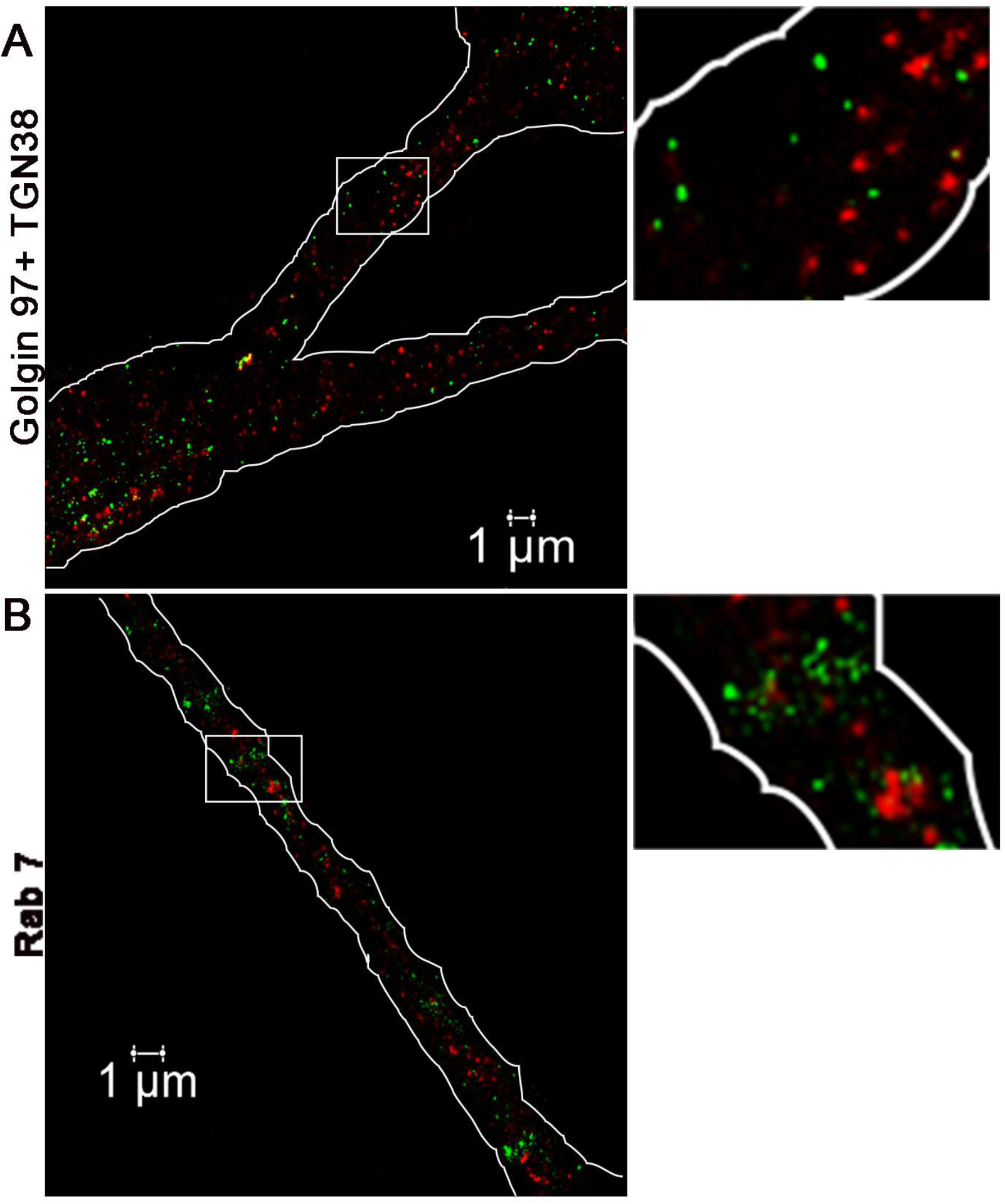
ATP7A containing vesicles do not colocalize with the TGN (Golgin 97+TGN38) markers and RAB7 in the neurites of the differentiated C-6 (glia) cells. ATP7A has been represented in green and the TGN markers and RAB7 in red. Image acquired by <u>S</u>timulated <u>E</u>mission <u>D</u>epletion (STED) microscopy.

**Supplementary figure 3:**
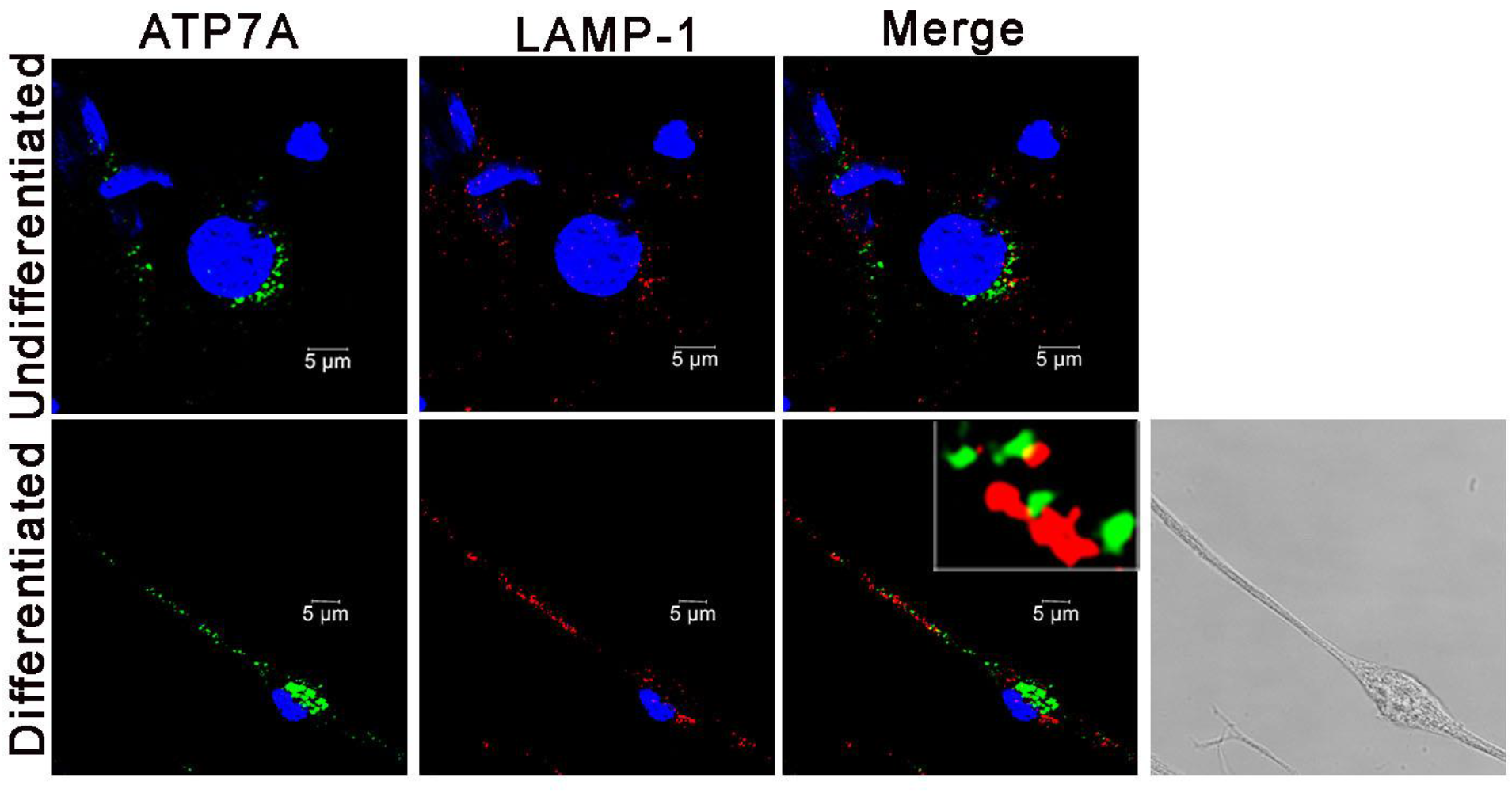
ATP7A containing vesicles do not colocalize with LAMP1 in the differentiated C-6 (glia) cells.

**Supplementary figure 4:**
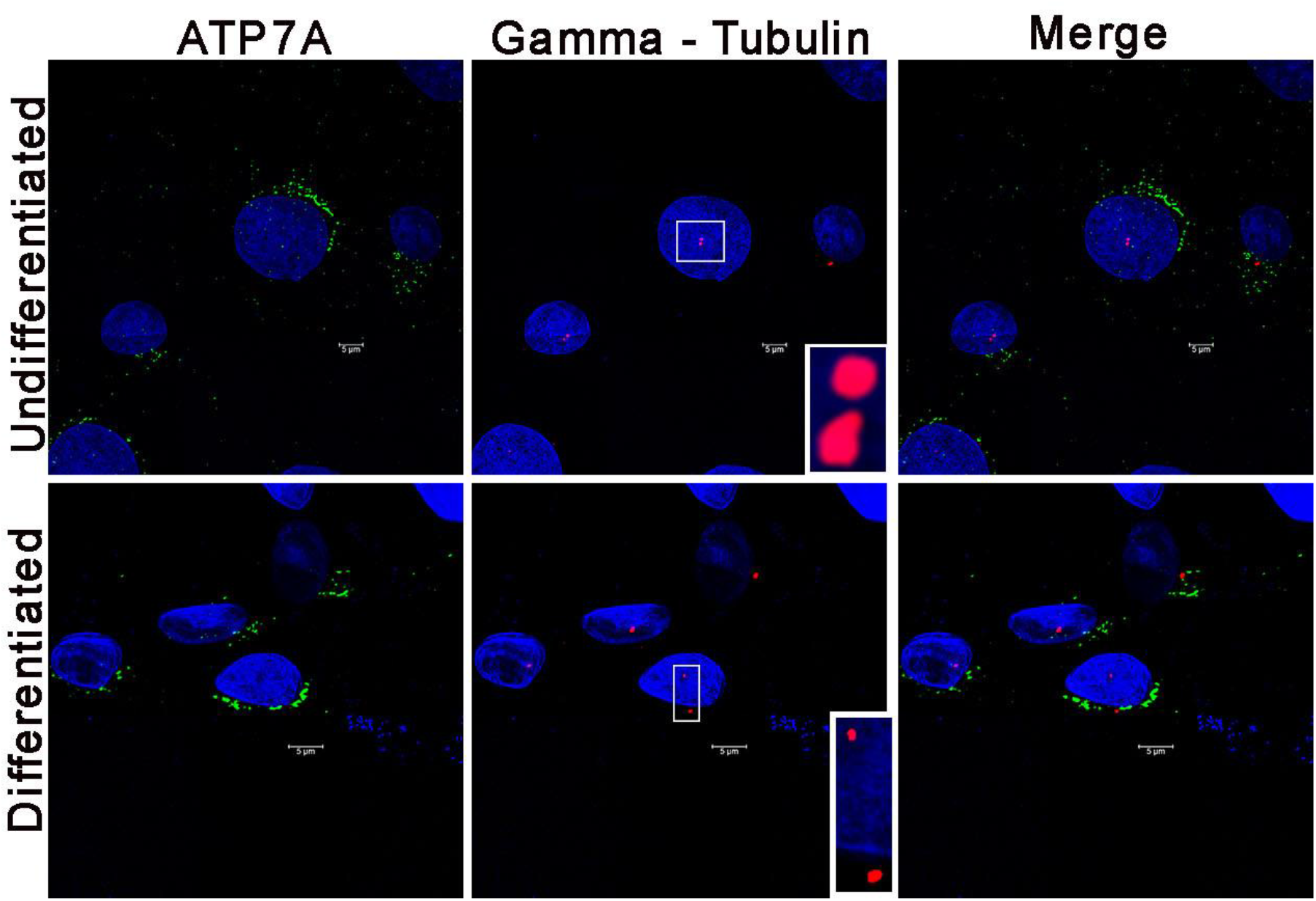
ATP7A containing vesicles do not colocalize with gamma-tubulin in the differentiated C-6 (glia) cells.

**Supplementary figure 5:**
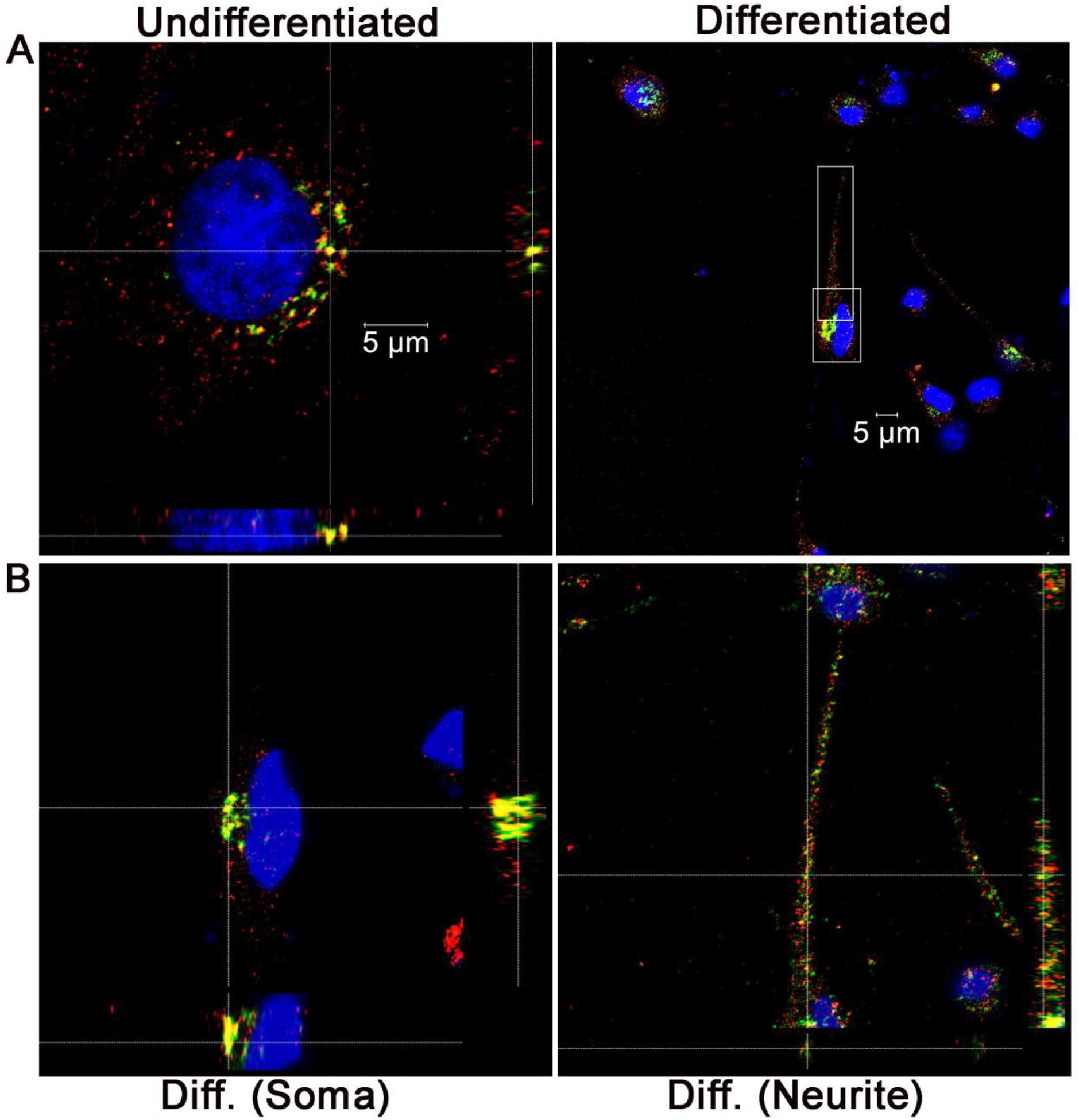
Colocalization of the ATP7A with RAB11 in the perinuclear (soma) region and along the neurites in the differentiated C-6 (glia) cells. A: Orthogonal (X/Y) images of ATP7A and RAB11 in the undifferentiated and differentiated C-6 cells has been represented. Region highlighted by boxes in the differentiated cell has been enlarged and represented in panel B. B – Orthogonal (X/Y) images of the enlarged perinuclear (soma) and neurite of a single cell, demonstrated in the panel A (right) and figure 4, have been represented.

**Supplementary figure 6:**
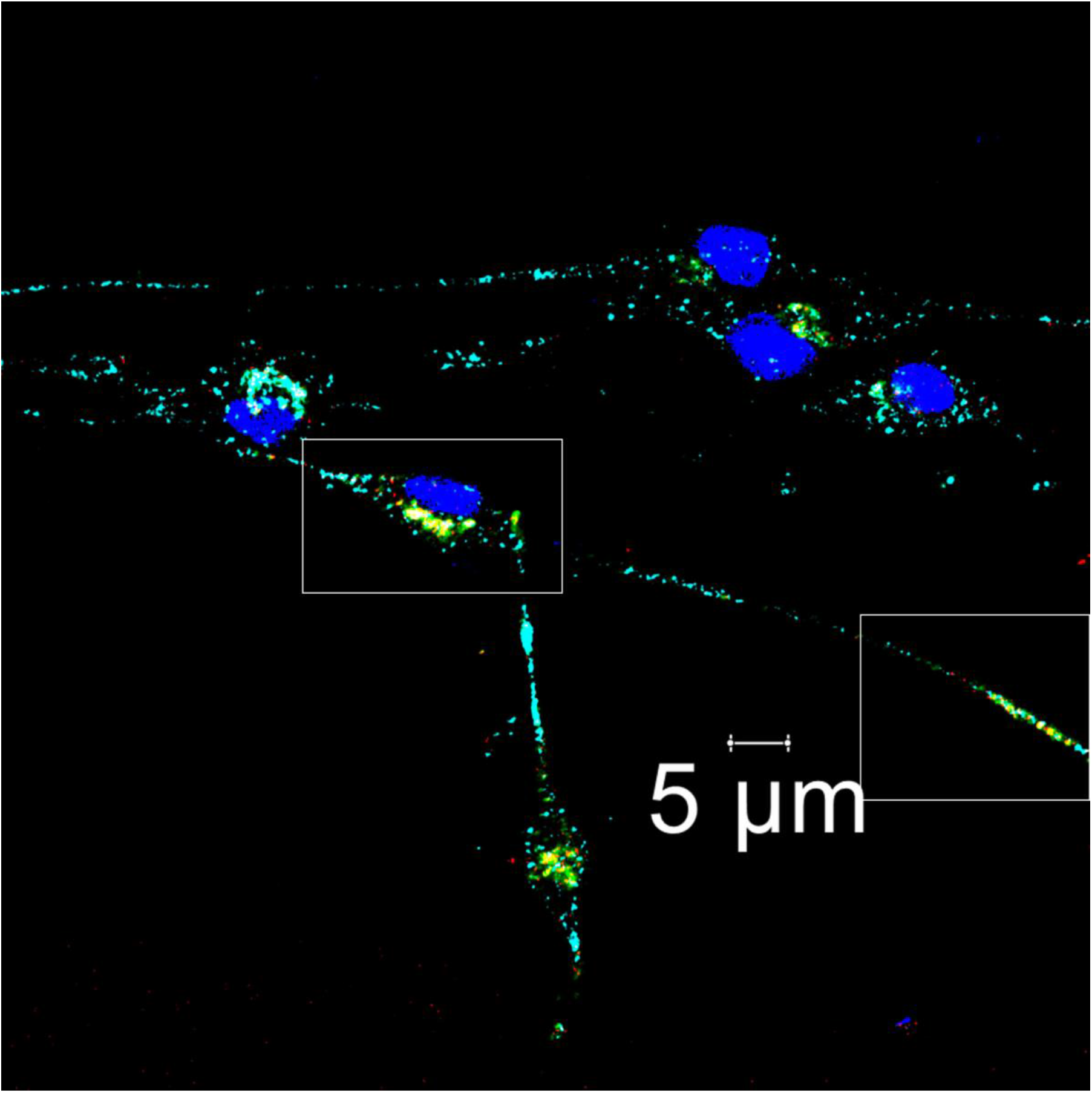
Distribution of ATP7A in TGN38+Golgin97 and RAB11 positive compartments in the differentiated C-6 (glia) cells. Magnified image of the highlighted regions has been represented in figure 6 and 7.

**Supplementary figure 7:**
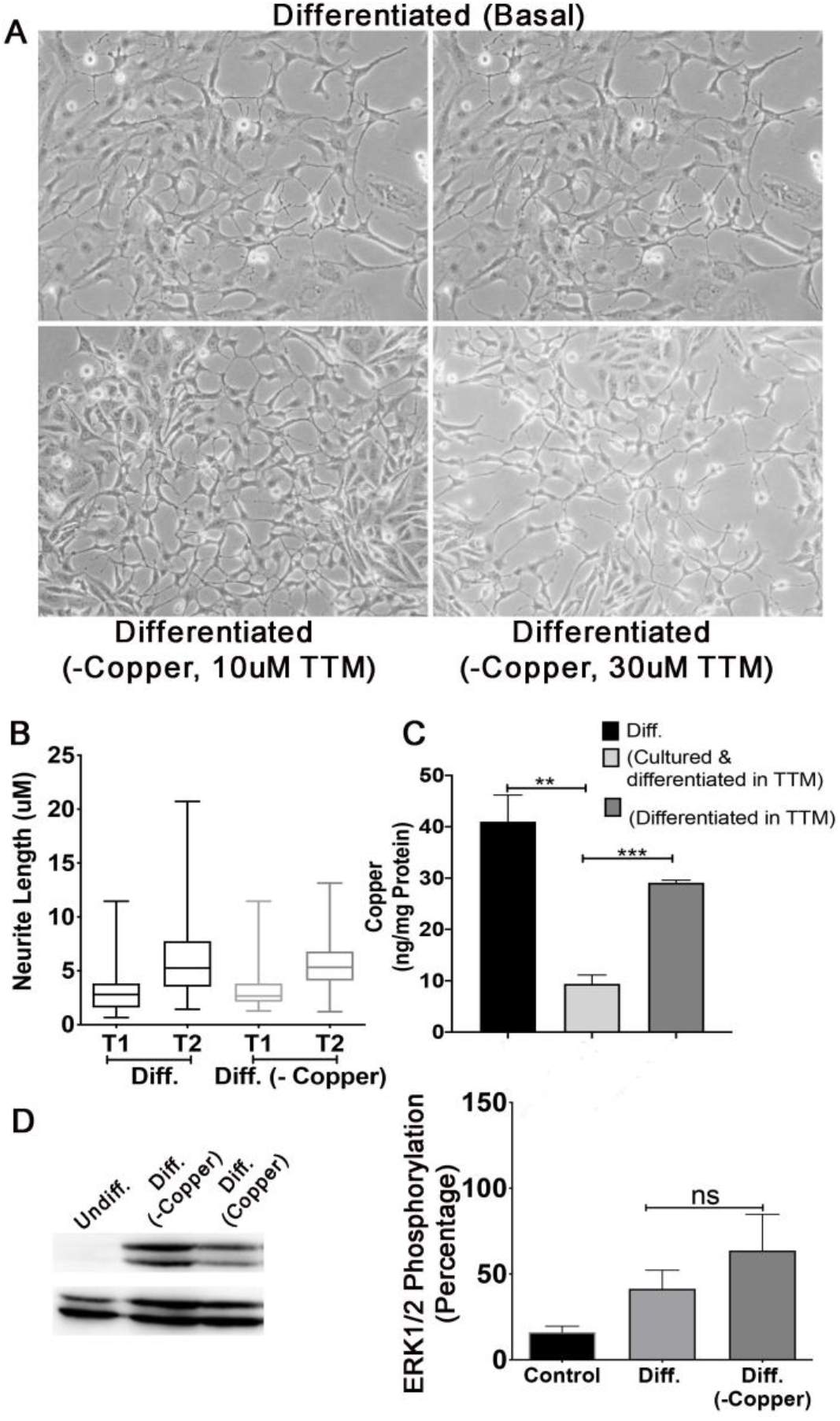
Copper chelation at the onset of the neuronal (PC-12) differentiation neither affects neurite outgrowth nor ERK1/2 phosphorylation. A - Bright field image of the PC12 cells differentiated in the presence and absence of TTM. B: Quantitation of the neurite length in the PC-12 cells differentiated in the presence and absence of TTM. C: Level of the intracellular copper in the PC-12 cells treated with the copper chelator TTM. D: Western Immunoblot (left) and quantitation (right) by densitometry of the phosphorylated ERK1/2 in the PC-12 cells under similar condition as mentioned.

**Supplementary figure 8:**
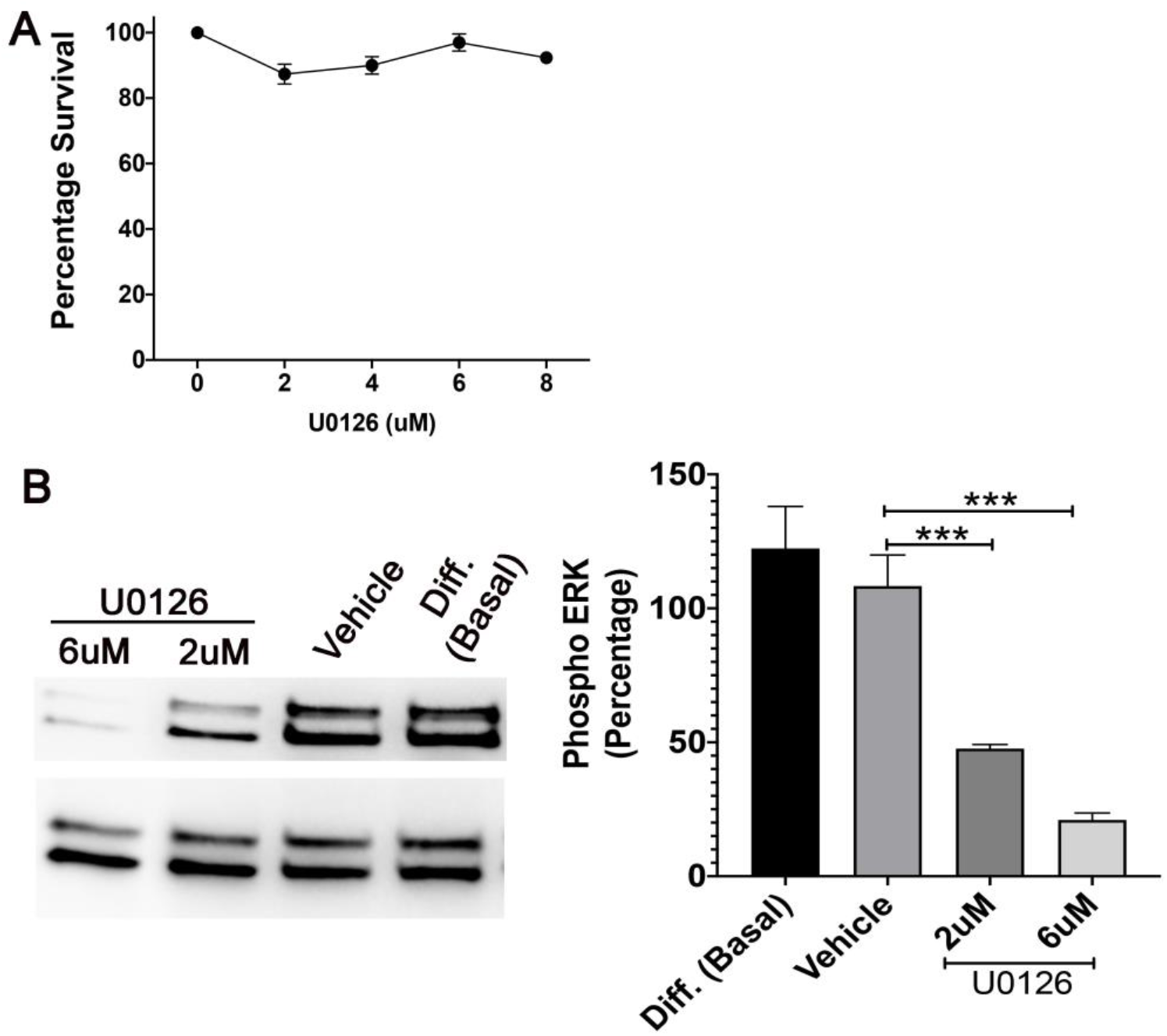
Inhibition of the ERK1/2 phosphorylation by U0126 in the PC-12 cells. A – Viability of the undifferentiated PC-12 cells in response to varying concentrations of U0126. B –. PC-12 cells are differentiated in presence of the varying concentrations of U0126. Western Immunoblot (left) and percentage of the Phospho ERK1/2 (right) calculated from the densitometric quantitation has been represented.

**Supplementary table 1:**
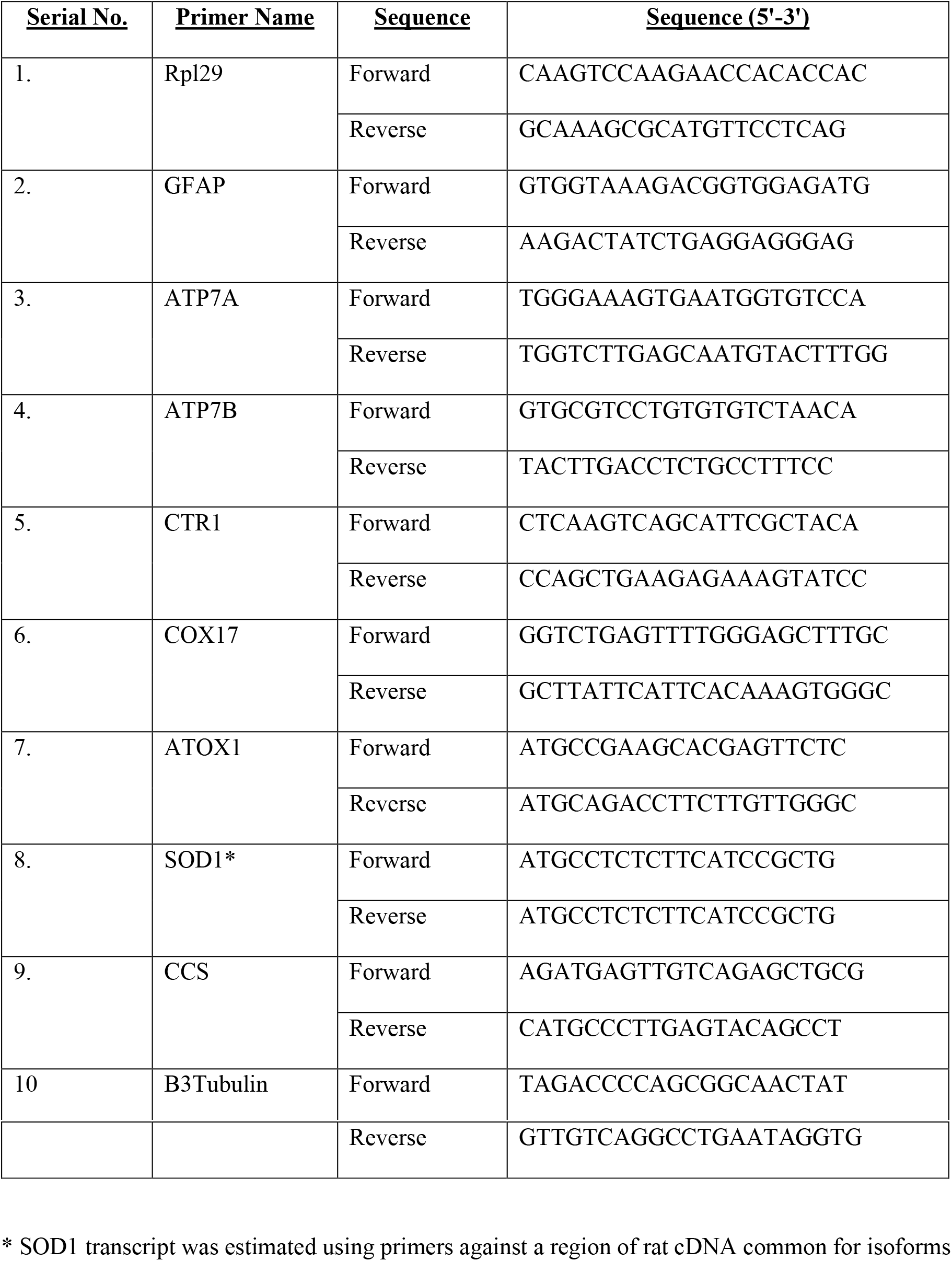
Primers used for amplification of mRNA for copper chaperones and transporters by real time PCR.

